# AtlasFold: Protein structure prediction with metagenomic-scale language models

**DOI:** 10.64898/2026.09.04.749352

**Authors:** Seonghwan Seo, Hyeongwoo Kim, Seokhyun Moon, Woo Youn Kim, Team KAIST

**Author notes:** Correspondence to Seonghwan Seo and Woo Youn Kim.

## Abstract

Protein language models (PLMs) trained on evolutionary sequences learn representations that encode protein structure, enabling direct structure prediction without multiple-sequence alignments (MSAs). Here we present the Atlas model family, an open and trainable system spanning protein language modeling, monomer folding, and protein-complex prediction. AtlasLM-3B is a 3B-scale language model trained with masked language modeling on approximately 1.56 billion sequences, including metagenomic data, and outperforms the similarly sized ESM2-3B in unsupervised contact prediction. Building on these representations, AtlasFold predicts all-atom protein structures and achieves state-of-the-art accuracy among PLM-based folding models. Fine-tuning AtlasFold for protein-complex prediction produces AtlasFold-Multimer (AtlasFold-M), whose antibody–antigen prediction performance is comparable to that of AlphaFold3 and ESMFold2. This protein-specific folding architecture enables fast, memory-efficient inference with AtlasFold and AtlasFold-M. By releasing the training code and data, stage checkpoints, and model weights under the MIT License, we provide a foundation for advancing PLM-based protein structure prediction.^1^

## 1 Introduction

Predicting how proteins fold and interact is central to understanding biological function. AlphaFold2 (Jumper et al., 2021) demonstrated high-accuracy prediction of individual protein structures by leveraging evolutionary information from multiple-sequence alignments (MSAs). Building on this advance, AlphaFold-Multimer (Evans et al., 2021), AlphaFold3 (Abramson et al., 2024), and co-folding models (Chai Discovery et al., 2024; Wohlwend et al., 2024; Zhang et al., 2026) extended structure prediction from individual proteins to biomolecular interactions.

Protein language models (PLMs) have emerged as an alternative to MSAs for protein structure prediction. Rao et al. (2020) showed that residue contacts are reflected in attention maps learned by PLMs through unsupervised language modeling. Extending this line of work, ESMFold (Lin et al., 2023) advanced PLM-based structure prediction from residue-level contacts to atomic-level protein structures directly from sequence, without an MSA search. More recently, Protenix-Mini (Gong et al., 2025) and ESMFold2 (Candido et al., 2026) have extended PLM-based structure prediction to molecular interactions. Together, these studies establish protein language modeling as a basis for predicting both monomer structures and protein interactions directly from sequence.

**Figure 1:**
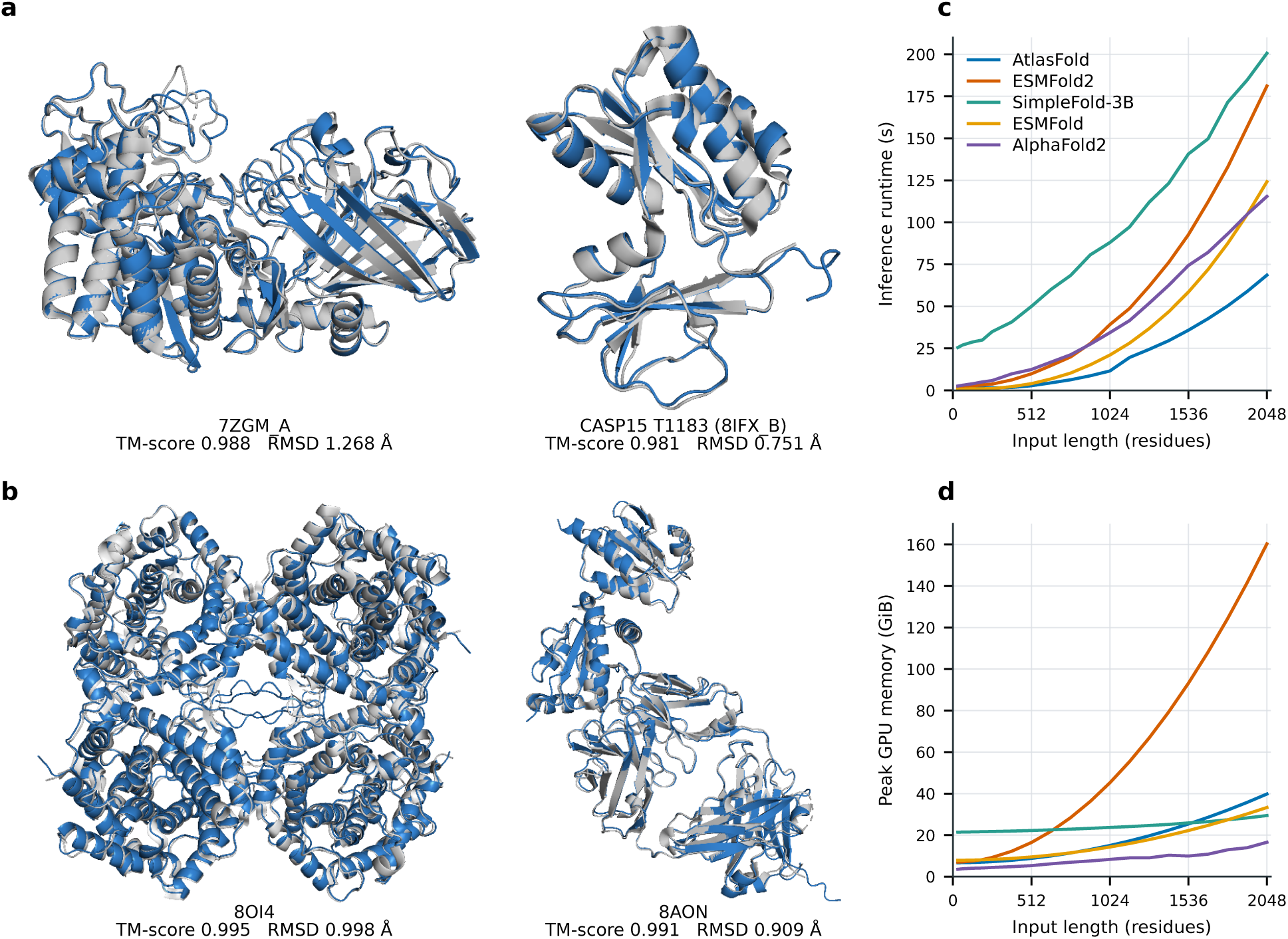
**(a–b)** Example predictions from AtlasFold **(a)** and AtlasFold-M **(b)**, with reference structures in gray and predictions in blue. **(a)** Monomer examples: 7ZGM_A and CASP15 target T1183 (8IFX_B). **(b)** Complex examples: the 8OI4 tetramer and the 8AON Fab complex. **(c–d)** Inference latency and peak GPU memory versus sequence length. Measurements use a single B200 GPU and exclude MSA-search time. AtlasFold and ESMFold2 generate five diffusion samples, SimpleFold-3B generates one sample, and AlphaFold2 uses one model. Inference settings and raw values are provided in Section D.

Here we introduce the Atlas model family, an open and trainable PLM-based system for protein folding and cofolding. At its foundation, AtlasLM is pretrained on a large corpus that includes metagenomic protein sequences. These sequences span broad evolutionary variation across protein families. AtlasLM-3B outperforms the similarly sized ESM2-3B in unsupervised long-range contact prediction, showing that its representations encode strong structural constraints.

Building on AtlasLM-3B, we develop AtlasFold to predict all-atom protein structures. AtlasFold is a state-of-the-art PLM-based folding model across CAMEO22, CASP14, and CASP15, while remaining competitive with MSA-based methods on CAMEO22. We further fine-tune AtlasFold on protein complexes for 25,000 optimizer steps to produce AtlasFold-Multimer (AtlasFold-M). On the FoldBench benchmark (Xu et al., 2025), AtlasFold-M achieves success rates (DockQ *>* 0.23) of 46.96% and 63.36% on the antibody–antigen and protein–protein subsets, respectively. AtlasFold and AtlasFold-M also provide fast, memory-efficient inference and support batching. We release the training code and data, stage checkpoints, and model weights under the MIT License.

## 2 Atlas model family

The Atlas model family comprises AtlasLM, AtlasFold, and AtlasFold-Multimer (AtlasFold-M). AtlasLM is pretrained with a masked-language-modeling objective on a large corpus that includes metagenomic protein sequences. AtlasFold combines representations from AtlasLM with a folding trunk and diffusion model to predict all-atom monomer structures, as illustrated in Fig. 3. Fine-tuning AtlasFold on protein complexes produces AtlasFold-M for multichain structure prediction. The following subsections summarize each Atlas model and its evaluation; detailed architecture, data, losses, and optimization schedules are provided in Sections A to C.

### 2.1 AtlasLM supports unsupervised contact prediction

AtlasLM-3B is a 3.06-billion-parameter transformer protein language model trained with masked language modeling. Its training collections contain approximately 1.56 billion sequences from UniRef (Suzek et al., 2007) and the metagenomic datasets MGnify (Mitchell et al., 2020) and MetaClust (Steinegger and Söding, 2018). Complete pretraining and model specifications are provided in Section A.

AtlasLM produces residue-level hidden states and pairwise attention maps. The hidden states describe each amino acid in sequence context, whereas the attention maps describe relationships between sequence positions. We assess whether these attention maps recover long-range contacts without structural supervision.

We evaluate AtlasLM-3B on partition 4 of the ESM structural superfamily split using the unsupervised attention-probe protocol of Rao et al. (2021) and report P@L and P@L/5 for long-range contacts. ESM models spanning 150 million to 15 billion parameters provide the comparison. At a similar parameter scale, AtlasLM-3B outperforms ESM2-3B on both P@L and P@L/5 (Fig. 2). It also outperforms ESM2-15B and the 300M and 600M ESMC models, while ESMC-6B achieves the highest precision on both metrics. Dataset filtering, contact definitions, probe fitting, and statistical procedures are provided in Section A.3.

**Figure 2:**
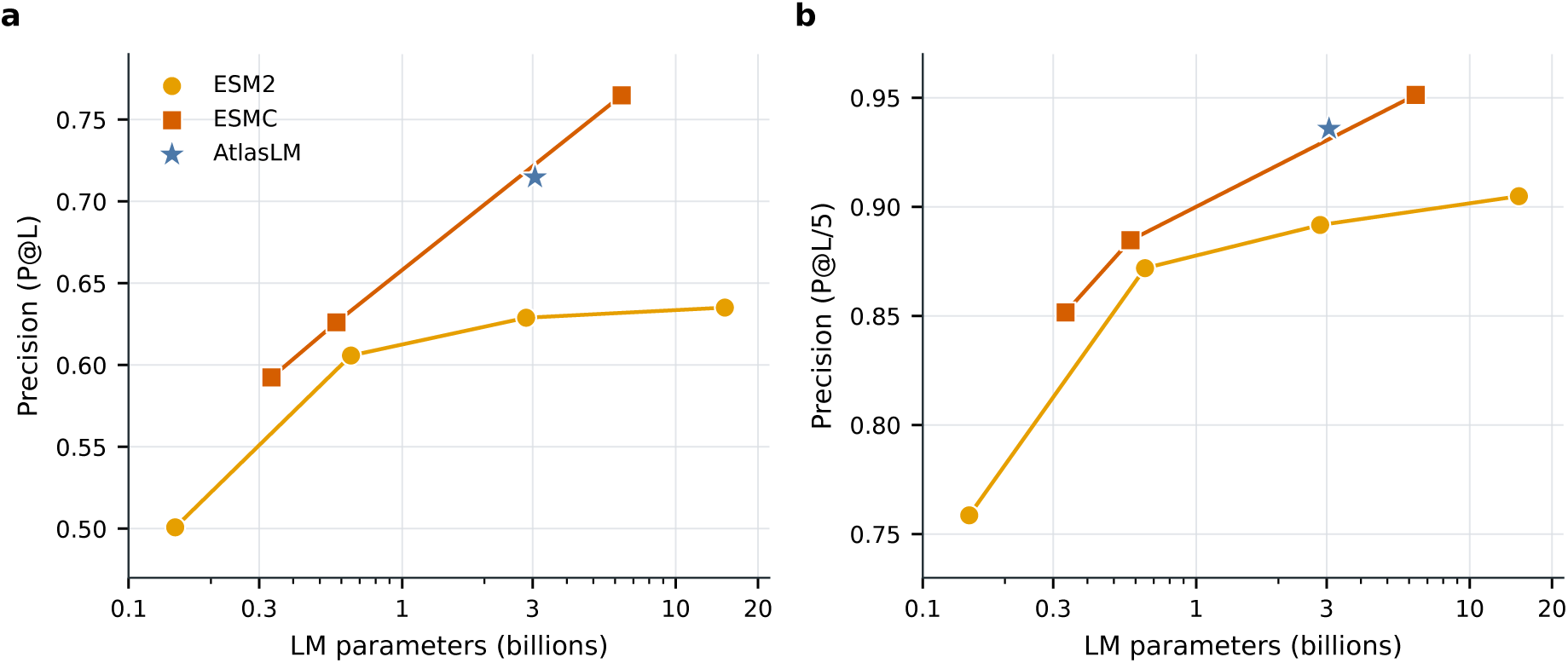
Unsupervised long-range contact precision for AtlasLM, ESM2, and ESMC on partition 4 of the ESM structural split. **(a)** P@L. **(b)** P@L/5. The detailed protocol is provided in Section A.3.

Together, these comparisons show that AtlasLM learns long-range pairwise structural information without structural supervision. AtlasLM-3B produces attention maps that are projected into the folding model’s pair representation.

### 2.2 AtlasFold predicts protein monomer structures

AtlasFold combines AtlasLM-3B representations with an AlphaFold3-derived folding trunk specialized for proteins (Abramson et al., 2024). At each recycling iteration, AtlasLM produces single and pair representations from an input sequence with 15% masking (Fig. 3a). The folding trunk combines these language-model representations with the recycled single and pair states, then updates both representations through a 48-block Pairformer.

**Figure 3:**
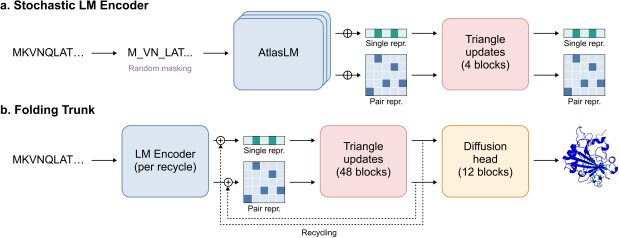
AtlasFold overview. **(a)** At each recycling iteration, AtlasLM recomputes single and pair representations from a newly masked input sequence. **(b)** The folding trunk combines the updated language-model representations with recycled state, refines them, and generates an all-atom structure with a diffusion head. Module details are provided in Section B.1.

Whereas AlphaFold3 uses a general molecular representation designed to accommodate heterogeneous inputs, including proteins, nucleic acids, modified residues, and ligands, AtlasFold restricts the prediction problem to protein monomers. This restricted scope allows every token to correspond to one amino acid, yielding a homogeneous protein representation. Within this protein-specific formulation, AtlasFold uses 4 recycles and a length-adaptive diffusion schedule^2^, rather than the 10 recycles and fixed 200-step schedule used by AlphaFold3. These architectural choices were motivated by our observation that diffusion sampling is the primary inference bottleneck for protein monomer prediction.

AtlasFold is trained on experimental PDB structures with a 2020-05-01 release-date cutoff and approximately 16 million AlphaFold2 predictions for MGnify sequences distributed through the OpenFold Portal (Ahdritz et al., 2024). Training proceeds in four stages with crop sizes of 256, 384, 512, and 640 residues for 45,000, 45,000, 8,000, and 2,000 optimizer steps, respectively. Additional architectural choices, dataset construction, and training procedures are provided in Section B.

#### 2.2.1 Monomer prediction accuracy

Monomer prediction is evaluated on CAMEO22 (*n* = 183) (Haas et al., 2018), CASP14 (*n* = 70), and CASP15 (*n* = 56). The comparison includes the MSA-based methods AlphaFold2 (Jumper et al., 2021), RoseTTAFold (Baek et al., 2021), and RoseTTAFold2 (Baek et al., 2023), as well as the PLM-based methods ESMFold (Lin et al., 2023), SimpleFold (Wang et al., 2026), and ESMFold2 (Candido et al., 2026). ESMFold2 is compared only on CASP15 because its training set overlaps CASP14 and its validation set overlaps CAMEO22.

To match the five structures generated by AlphaFold2 from its five models, AtlasFold and ESMFold2 generate one diffusion sample for each of five random seeds (1–5), while SimpleFold generates five diffusion samples. For methods with multiple candidates, the structure with the highest average pLDDT is selected; the evaluation protocol and full inference settings are described in Section B.3. Across CAMEO22, CASP14, and CASP15, AtlasFold is a state-of-the-art PLM-based folding model (Table 1). AtlasFold remains competitive with MSA-based methods on CAMEO22, while AlphaFold2 is stronger on CASP14 and CASP15.

**Table 1:** Monomer structure prediction on CAMEO22, CASP14, and CASP15. Values are mean / median; bold indicates the best result within each MSA-based or PLM-based group for each statistic.

| Type | Model | TM-score ↑ | GDT-TS ↑ | IDDT ↑ | IDDT-Cα ↑ | RMSD ↓ |
| --- | --- | --- | --- | --- | --- | --- |
| *CAMEO22* |  |  |  |  |  |  |
| MSA-based | RoseTTAFold | 0.780 / 0.860 | 0.715 / 0.775 | 0.575 / 0.605 | 0.798 / 0.827 | 5.721 / 2.864 |
|  | RoseTTAFold2 | 0.864 / 0.947 | 0.844 / 0.900 | 0.751 / 0.794 | 0.891 / 0.923 | 3.513 / 1.728 |
|  | AlphaFold2 | **0.879 / 0.955** | **0.863 / 0.914** | **0.826 / 0.869** | **0.904 / 0.932** | **3.178 / 1.574** |
| PLM-based | ESMFold | 0.853 / 0.933 | 0.826 / 0.875 | 0.791 / 0.832 | 0.871 / 0.906 | 3.995 / 2.018 |
|  | SimpleFold-3B | 0.835 / 0.911 | 0.801 / 0.854 | 0.776 / 0.807 | 0.855 / 0.889 | 4.234 / 2.183 |
|  | AtlasFold | **0.865 / 0.945** | **0.847 / 0.907** | **0.838 / 0.880** | **0.898 / 0.936** | **3.538 / 1.909** |
| *CASP14* |  |  |  |  |  |  |
| MSA-based | RoseTTAFold | 0.654 / 0.678 | 0.562 / 0.572 | 0.464 / 0.456 | 0.705 / 0.723 | 9.676 / 6.420 |
|  | RoseTTAFold2 | 0.802 / 0.881 | 0.744 / 0.815 | 0.670 / 0.706 | 0.832 / 0.877 | 6.614 / 3.167 |
|  | AlphaFold2 | **0.844 / 0.893** | **0.784 / 0.842** | **0.778 / 0.803** | **0.865 / 0.893** | **4.481 / 2.881** |
| PLM-based | ESMFold | 0.702 / 0.790 | 0.621 / 0.713 | 0.635 / 0.700 | 0.722 / 0.797 | 8.652 / 4.069 |
|  | SimpleFold-3B | 0.716 / 0.788 | 0.638 / 0.705 | 0.666 / 0.697 | 0.748 / 0.829 | 7.969 / 4.048 |
|  | AtlasFold | **0.732 / 0.804** | **0.650 / 0.724** | **0.683 / 0.748** | **0.751 / 0.839** | **7.299 / 3.988** |
| *CASP15* |  |  |  |  |  |  |
| MSA-based | RoseTTAFold | 0.639 / 0.685 | 0.550 / 0.554 | 0.636 / 0.700 | 0.721 / 0.790 | 13.961 / 6.740 |
|  | RoseTTAFold2 | 0.724 / 0.843 | 0.668 / 0.725 | 0.644 / 0.746 | 0.803 / 0.877 | 14.302 / 4.500 |
|  | AlphaFold2 | **0.751 / 0.854** | **0.702 / 0.747** | **0.759 / 0.829** | **0.842 / 0.897** | **9.759 / 4.870** |
| PLM-based | ESMFold | 0.671 / 0.759 | 0.608 / 0.667 | 0.671 / 0.774 | 0.755 / 0.835 | 14.259 / 8.323 |
|  | ESMFold2 | 0.695 / 0.734 | **0.645 / 0.712** | **0.726 / 0.827** | **0.790 / 0.892** | 14.499 / **5.755** |
|  | SimpleFold-3B | 0.653 / 0.694 | 0.575 / 0.573 | 0.637 / 0.700 | 0.728 / 0.804 | 15.802 / 8.380 |
|  | AtlasFold | **0.701 / 0.789** | **0.645 / 0.711** | 0.725 / 0.812 | **0.790 / 0.885** | **10.307 / 7.952** |

#### 2.2.2 Confidence evaluation

Following AlphaFold2 (Jumper et al., 2021), AtlasFold predicts per-residue pLDDT as an estimate of local structural accuracy and pTM as an estimate of global fold accuracy. Per-residue pLDDT identifies locally reliable regions, its mean over a predicted structure provides a score for selecting among sampled structures, and pTM summarizes confidence in the complete fold.

We evaluate confidence on 10,000 randomly selected monomer structures released after 2023-01-01, excluding targets with a structural-training-set match at least 40% sequence identity and more than 80% target coverage. This filter applies to the structural-training set but not to the AtlasLM pretraining corpus, which may contain related sequences.

Mean pLDDT over experimentally resolved residues tracks lDDT-*C_α_* on the same residues, with Pearson *r* = 0.829 and Spearman *ρ* = 0.765, while resolved-region pTM tracks TM-score with *r* = 0.909 and *ρ* = 0.894 (Fig. 4b). Together, these comparisons evaluate the confidence outputs for one prediction per target across the holdout.

**Figure 4:**
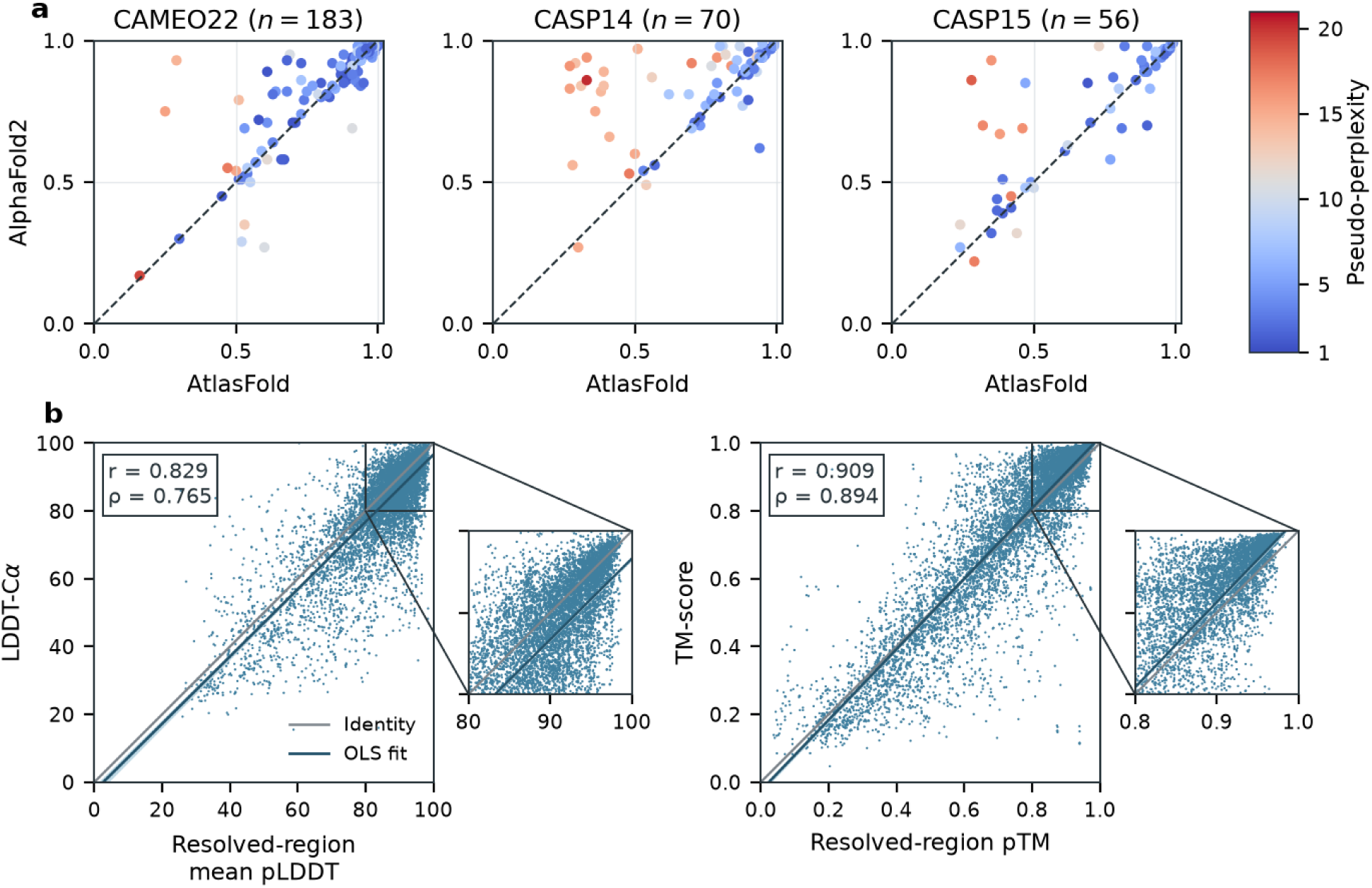
Relationship between AtlasLM pseudo-perplexity, structure accuracy, and AtlasFold confidence. **(a)** AtlasFold and AlphaFold2 TM-scores on CAMEO22, CASP14, and CASP15, colored by AtlasLM pseudo-perplexity. **(b)** Mean pLDDT over experimentally resolved residues versus lDDT-*C_α_* on the same residues, and resolved-region pTM versus TM-score, on the 10,000-target confidence holdout. Here, *r* and *ρ* denote Pearson and Spearman correlations, respectively.

AtlasLM pseudo-perplexity provides a separate, sequence-level view of prediction difficulty (Fig. 4a). AtlasFold and AlphaFold2 have similar TM-scores on many low-pseudo-perplexity targets, while several of the largest differences in favor of AlphaFold2 occur on proteins with high pseudo-perplexity, particularly in CASP14 and CASP15. This association is descriptive and strongest on CASP14; it is weaker on CAMEO22 and CASP15 and does not by itself establish why the errors occur.

### 2.3 Fine-tuning AtlasFold for protein-complex prediction

We next investigate whether AtlasFold can be adapted for protein-complex prediction. We fine-tune AtlasFold on protein complexes for 25,000 optimizer steps to produce AtlasFold-Multimer (AtlasFold-M). AtlasFold-M retains the folding trunk of AtlasFold while adding a template module and an AlphaFold3-style confidence module (Abramson et al., 2024). Detailed architectural changes, training data, and objectives are provided in Section C. Although a template module is introduced, all experiments reported in this study were conducted without templates.

**Figure 5:**
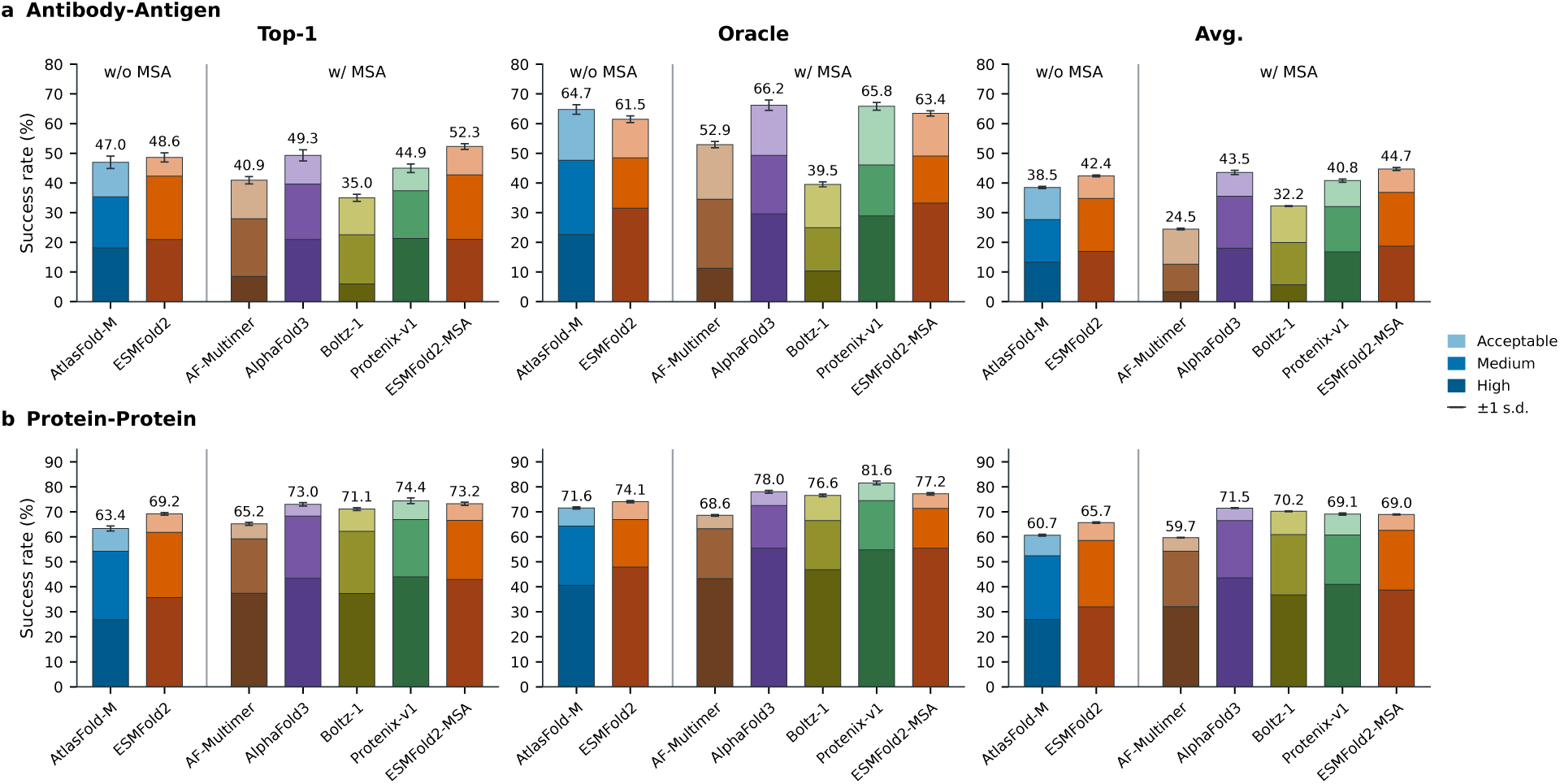
Protein-complex prediction on the **(a)** antibody–antigen and **(b)** protein–protein subsets of FoldBench. Bars show the fractions above the indicated DockQ thresholds. Top-1 uses model confidence, Oracle selects the highest-DockQ candidate, and Avg. averages across generated candidates. Results are means across all five-seed subsets drawn from 10 seeds, and error bars denote the corresponding standard deviations. Detailed protocols and raw values are provided in Section C.3.2.

**Figure 6:**
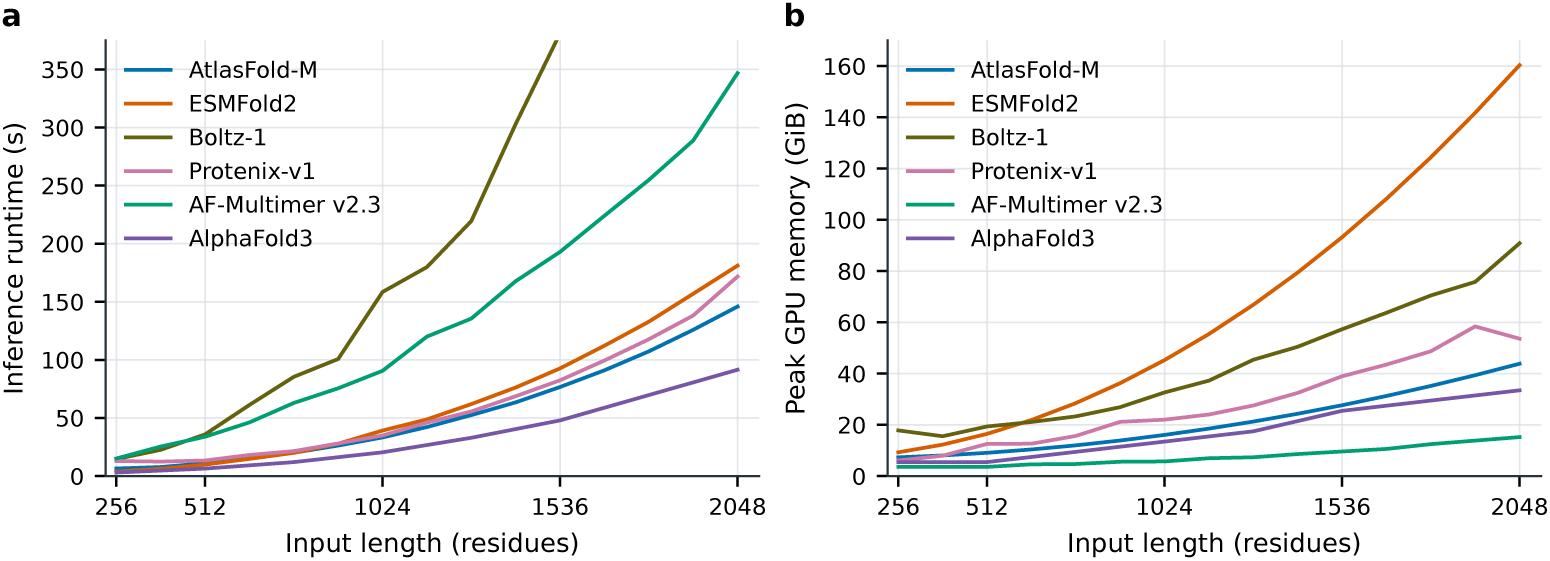
Inference runtime **(a)**, and peak GPU memory **(b)**, on one B200 GPU. The reported inference settings differ across models and are detailed in the supplementary information. Inference settings and raw values are provided in Section D.

#### 2.3.1 Protein-complex prediction

We evaluate AtlasFold-M on the protein–protein (P–P) and antibody–antigen (Ab–Ag) subsets of FoldBench (Xu et al., 2025). The evaluation covers all 172 Ab–Ag targets and 278 of the 279 P–P targets, excluding 8GQP, a D-protein binder system. AtlasFold-M uses 10 recycles and 200 diffusion steps. Additional protocol and baseline details are provided in Section C.

With confidence-based selection, AtlasFold-M achieves DockQ success rates (*>* 0.23) of 63.36% on P–P and 46.96% on Ab–Ag (Fig. 5). Its Ab–Ag performance is comparable to that of state-of-the-art methods, including AlphaFold3 and ESMFold2. On the P–P subset, AtlasFold-M performs below the other co-folding models evaluated here, which may reflect the smaller size of AtlasLM-3B and the limited scale of multimer fine-tuning. Complete results for DockQ, Fnat, iRMSD, LRMSD, and lDDT are provided in Section C.3.2.

**Figure 7:**
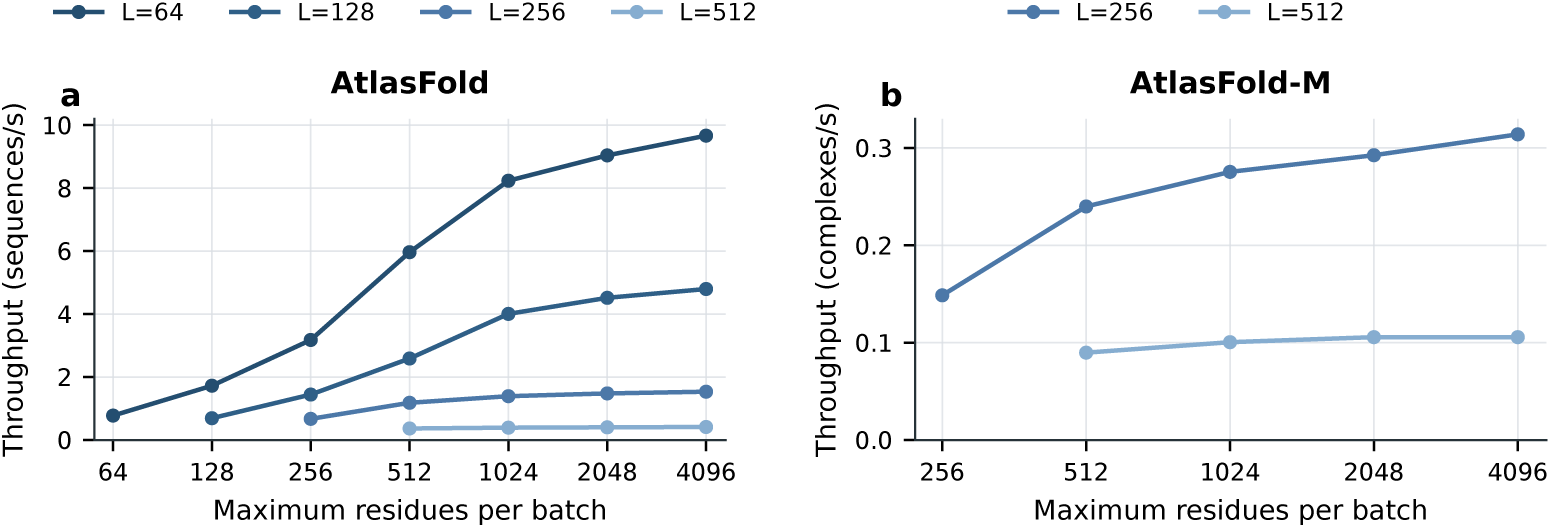
Throughput of batched inference for **(a)** AtlasFold monomer and **(b)** AtlasFold-M at the indicated sequence or complex lengths. The number of diffusion samples is five. The 64-residue point in **(a)** uses the corresponding single-run measurement. Raw values are provided in Section D.

### 2.4 Fast and batched inference for monomers and complexes

We profiled monomer and complex inference on a single NVIDIA B200 GPU and measured inference runtime, peak GPU memory, and batched throughput (Figs. 1c,d, 6, and 7). Complete inference settings and measurements are provided in Section D.

AtlasFold provides fast monomer prediction across sequence lengths from 32 to 2,048 residues. At 1,024 residues, AtlasFold generates five structures in 11.57 s, compared with 34.38 s for one AlphaFold2 model, 39.17 s for five ESMFold2 structures, and 88.01 s for one SimpleFold-3B structure. AtlasFold also uses less peak memory than ESMFold2 throughout the measured range, requiring 39.81 GiB rather than 160.32 GiB at 2,048 residues.

AtlasFold-M remains efficient for complexes containing up to 2,048 total residues. Using five samples, 10 recycles, and 200 diffusion steps, AtlasFold-M requires 33.24 s and 16.05 GiB at 1,024 total residues, compared with 34.96 s and 22.00 GiB for Protenix-v1, 39.17 s and 45.24 GiB for ESMFold2, 90.59 s and 5.70 GiB for AlphaFold-Multimer v2.3, and 158.44 s and 32.63 GiB for Boltz-1. Its peak memory is lower than that of ESMFold2 across all measured complex lengths.

Batching further improves throughput when many targets are predicted. For 64-residue monomers, batched inference raises AtlasFold throughput from 0.775 to 9.663 sequences/s, a 12.5-fold gain. For 256-residue complexes, batched inference raises AtlasFold-M throughput from 0.149 to 0.314 complexes/s, a 2.1-fold gain.

### 2.5 Hallucinations in disordered regions

The AlphaFold3 study (Abramson et al., 2024) describes spurious structural order in disordered regions as hallucination. Although these regions are typically assigned low confidence, their generated coordinates can lack the extended, ribbon-like appearance produced by AlphaFold2 and AlphaFold-Multimer v2.3. AlphaFold3 mitigates this behavior through cross-distillation from AlphaFold-Multimer predictions and a ranking term that favors greater solvent-accessible surface area. We examine the same two examples used in the AlphaFold3 analysis: the CAID2 (Conte et al., 2023) target DP02376 and the 7F60 nuclear pore complex.

For DP02376, AlphaFold3 and ESMFold2 produce extended low-confidence chains, while AtlasFold retains extended trajectories with more packing around the predicted structured regions (Fig. 8a). ESMFold also yields spurious structural order, with low-confidence residues packed around the predicted domains despite using an invariant point attention structure module rather than a diffusion coordinate generator (Lin et al., 2023). These packed conformations resemble those produced by diffusion models, suggesting that this hallucination is not specific to diffusion-based coordinate generation.

**Figure 8:**
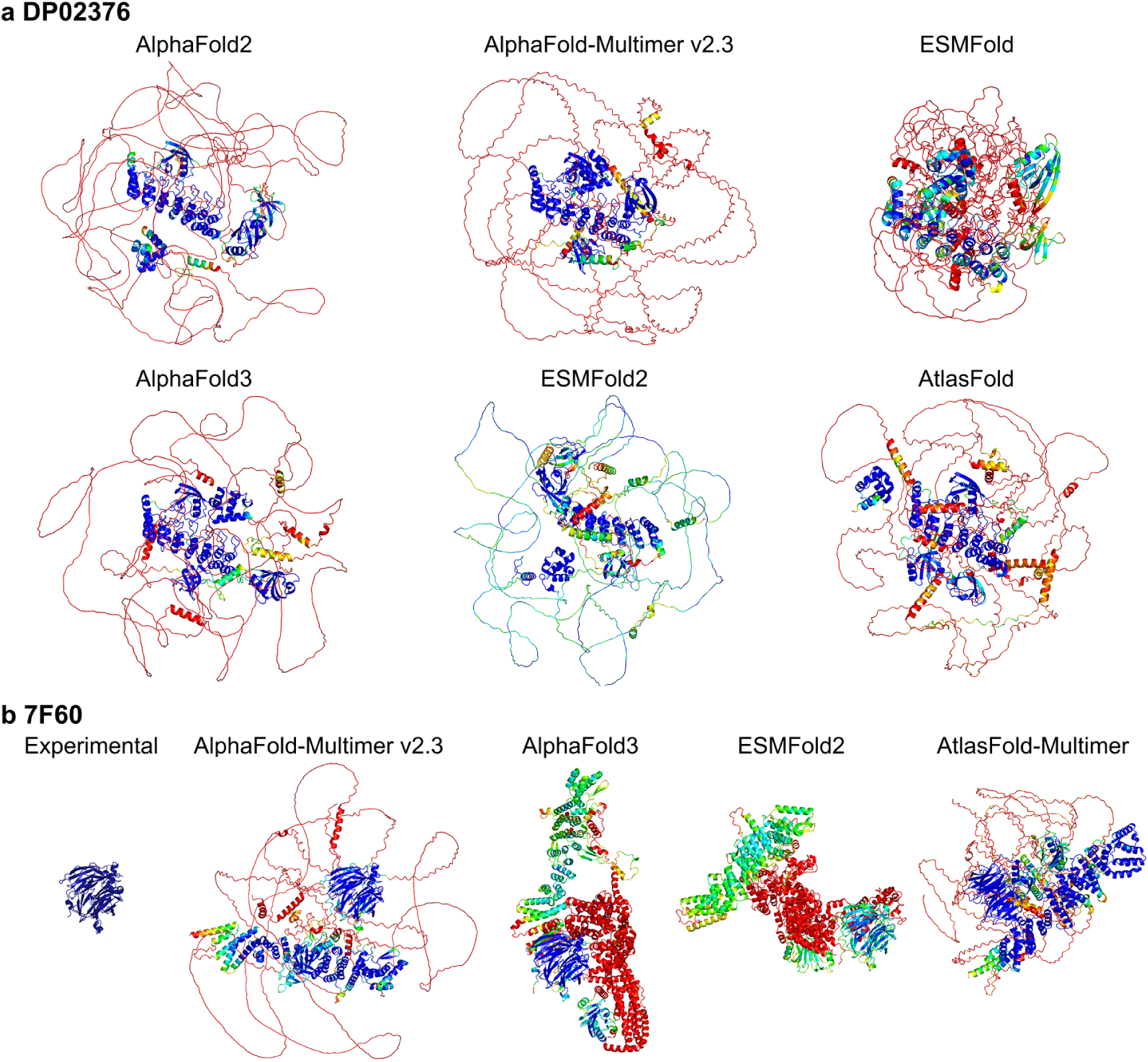
Qualitative comparison of low-confidence disordered-region predictions. **(a)** Predictions for the CAID2 target DP02376. **(b)** Resolved experimental structure and predictions for the 7F60 nuclear pore complex, whose experimental structure omits 1,854 unresolved residues. Predictions are colored by pLDDT, and the highest-ranked output is shown.

For 7F60, spurious structural order appears in all five AlphaFold3 and ESMFold2 samples and in one of the five AtlasFold-M samples generated with seed 1. The highest-ranked AtlasFold-M sample shown in Fig. 8b is not the ordered sample; it retains more extended trajectories that remain packed around the predicted structured core. The resemblance of AtlasFold-M to AlphaFold-Multimer v2.3 is consistent with the AlphaFold-Multimer augmentation used during multimer fine-tuning. In these examples, low-confidence packing differs across models and cannot be attributed to the coordinate-generation architecture alone.

## 3 Design and training observations

Several choices across AtlasLM and AtlasFold emerged from observations made during development.

AtlasLM was developed before the release of ESMC and ESMFold2, and its data preparation and training schedule instead followed ESM3 (Hayes et al., 2025). Because reconstructing the two-billion-cluster JGI collection later reported for ESMC was impractical, we used the smaller MetaClust collection in its place. Long-range contact precision continued to improve across checkpoints, but at 1.5 million updates AtlasLM remained below the performance reported for ESMC, likely reflecting differences in dataset scale and training settings (Candido et al., 2026). We therefore continued training for another 0.5 million updates, reaching 2.0 million updates in total (Fig. 9).

**Figure 9:**
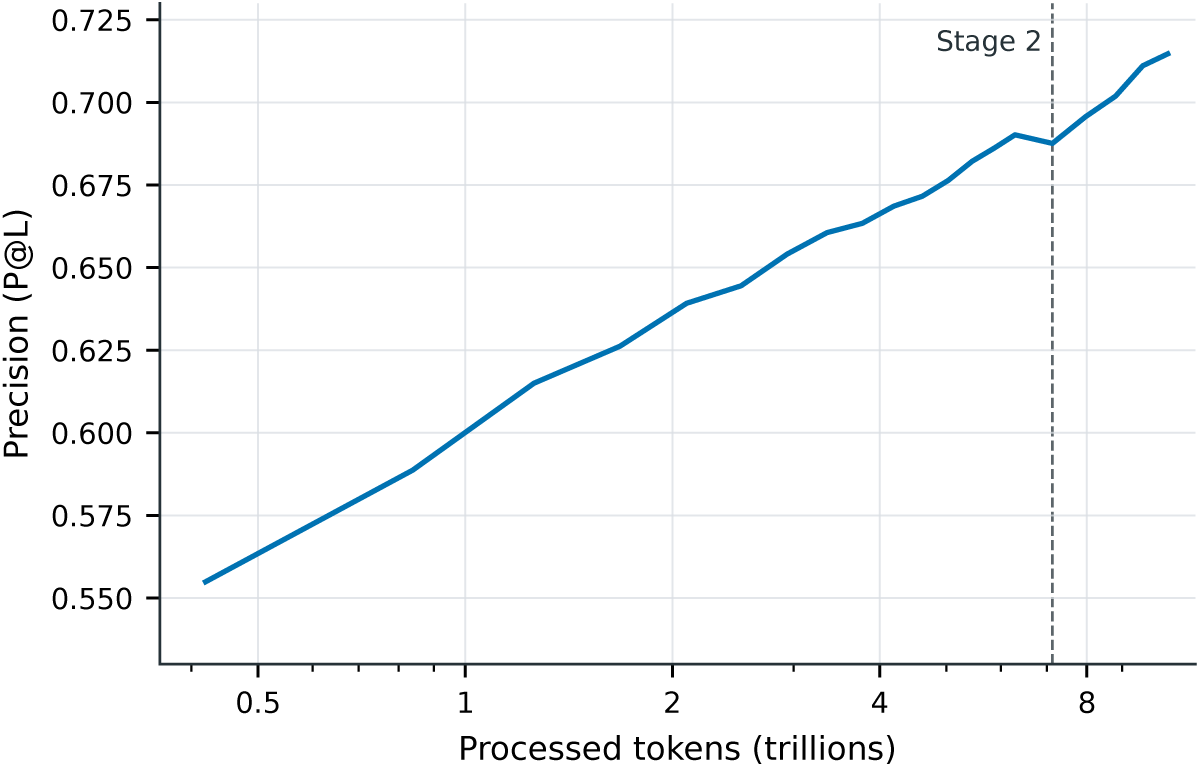
AtlasLM-3B P@L across training checkpoints. The dashed line marks the transition to stage 2.

During development, we evaluated ESMFold2 on CASP14, although CASP14 is included in its training set. ESMFold2 reached a mean TM-score of 0.736 and a mean GDT-TS of 0.661, only modestly above AtlasFold’s 0.732 and 0.650 and still well below AlphaFold2’s 0.844 and 0.784. This result suggests that the limitation of PLM-based folding extends beyond generalization to unseen targets: training-set exposure alone did not close the gap to an MSA-based model. For monomer folding, representations learned through masked language modeling therefore appear less directly aligned with three-dimensional structure prediction than the explicit coevolutionary constraints extracted from MSAs. ESMFold2, which uses ESMC-6B, outperforms AtlasFold-M on both FoldBench subsets, reaching confidence-ranked acceptable-or-better success rates of 69.22% versus 63.36% on protein–protein targets and 48.60% versus 46.96% on antibody–antigen targets. These co-folding results motivate exploring larger PLM backbones for AtlasFold-M.

Inspired by ESMFold2, we initially removed triangle attention from the folding trunk. Despite changes to the optimizer and other training settings, this configuration repeatedly encountered gradient explosions around 30,000 updates. We therefore restored triangle attention and reduced the pair-representation width in the final architecture. We also tested AdamW following ESMFold2 and compared it with Adam, as used in AlphaFold2 (Jumper et al., 2021) and AlphaFold3 (Abramson et al., 2024). In our development runs, Adam reduced the training loss more rapidly and reached comparable validation accuracy in fewer updates than AdamW, so we used Adam for the final structure-training runs.

AlphaFold3 applies EDM sampling (Karras et al., 2022) to general biomolecular complexes. Because AtlasFold targets protein monomers, we tested whether the AlphaFold3 sampling parameters could be adjusted for this narrower setting. Reducing the initial noise level had little effect on overall prediction quality and increased the tendency toward packed conformations. Although the released model uses 100 denoising steps for sequences longer than 1,024 residues, a fixed 25-step schedule was sufficient for most targets. For some targets with low predicted TM-score, however, a small number of residues remained poorly resolved at the atomic level. We therefore adopted 20 denoising steps for sequences of up to 512 residues, 30 steps for sequences of 513–1,024 residues, and 100 steps for longer sequences.

The confidence module required a separate revision. We initially used an AlphaFold3-style shared module for local and pairwise confidence prediction. Adding PAE supervision degraded pLDDT-based sample ranking. We therefore separated the local and pairwise confidence paths, reinitialized the confidence module, and retrained it from scratch after stage 4.

During AtlasFold-M development, we incorporated AlphaFold-Multimer predictions for homodimers and heterodimers from an expanded version of the AlphaFold Database (AFDB) (Han et al., 2026). We tested sampling rates of 8% for homodimers and 2% for heterodimers during stage 2 training and observed meaningful improvements in protein–protein complex prediction. However, performance decreased on antibody–antigen targets, so we did not use the augmented-data configuration in the final AtlasFold-M release. Limited computational resources prevented us from exploring additional sampling ratios, but these results suggest that appropriately balanced synthetic structures predicted by AlphaFold-Multimer may benefit future co-folding training.

## 4 Conclusion

The Atlas family establishes an open and trainable path from metagenomic-scale protein language modeling to all-atom monomer and protein-complex prediction. AtlasLM-3B learns long-range structural constraints directly from evolutionary sequence data and outperforms the similarly sized ESM2-3B in unsupervised contact prediction. AtlasFold transfers these representations to a protein-specific folding architecture and achieves state-of-the-art accuracy among PLM-based monomer models across CAMEO22, CASP14, and CASP15. AtlasFold-M extends this framework to protein complexes and achieves competitive performance on the protein–protein and antibody–antigen subsets of FoldBench. Specializing the folding architecture to proteins further enables fast, memory-efficient inference and batching for both monomer and complex prediction.

For monomer folding, ESMFold2 performs similarly to AtlasFold on CASP14 despite using ESMC-6B and including CASP14 in its training set. This observation suggests that the limitations of PLM-based folding extend beyond generalization to unseen targets and that PLM scaling alone may not close the gap to MSA-based models. For co-folding, however, ESMFold2 outperforms AtlasFold-M on both the protein–protein and antibody–antigen subsets of FoldBench, suggesting that PLM scaling may remain meaningful for modeling inter-chain interactions.

The three Atlas models point to complementary directions for future work. AtlasLM-3B combines strong structural representations with a practical 3B-parameter scale, making it a useful backbone for downstream protein models operating at the residue, sequence, or structure level. For AtlasFold, a central direction is to align PLM-derived features more closely with folding, for example through folding-aware pretraining objectives, improved pair representations, or structure-guided adaptation that preserves the efficiency of single-sequence inference. For AtlasFold-M, the results demonstrate that a pretrained monomer folding model can provide a practical starting point for co-folding: 25,000 additional optimizer steps are sufficient to reach competitive antibody–antigen performance. By avoiding the need to train the language and folding components from scratch, this transfer can lower the cost of developing and studying co-folding models and facilitate experiments with larger language-model backbones and larger, more diverse complex training sets. The present framework remains limited to proteins and does not support ligands, DNA, or RNA; this limitation will be addressed in our follow-up work, K-Fold.

Beyond the final models, we release the training code and data, stage checkpoints, and model weights under the MIT License. Alongside these resources, we report practical observations from the development of AtlasLM, AtlasFold, and AtlasFold-M, spanning pretraining scale, folding architecture, optimization, sampling, confidence modeling, and complex fine-tuning. By sharing both the resources and the observations that shaped them, we hope to support an open scientific community in which protein structure models can be understood, questioned, and extended.

## Acknowledgments

This work was developed as part of the K-Fold initiative supported by the Ministry of Science and ICT (MSIT), Republic of Korea, through the National IT Industry Promotion Agency (NIPA) (Grant No. PJT-26-100009).

## Appendix

### A AtlasLM training and contact evaluation

#### A.1 Architecture

AtlasLM uses the pre-normalization Transformer architecture of ESMC (Candido et al., 2026). Each block applies rotary-position-encoded multi-head self-attention with query/key layer normalization, followed by a SwiGLU feed-forward network; both sublayers use bias-free linear projections and residual connections. A final layer normalization and a two-layer masked-token regression head produce amino-acid logits. Table 2 places AtlasLM-3B beside the ESM-2 (Lin et al., 2023) and ESMC (Candido et al., 2026) models included in the main-text contact experiment.

**Table 2:**
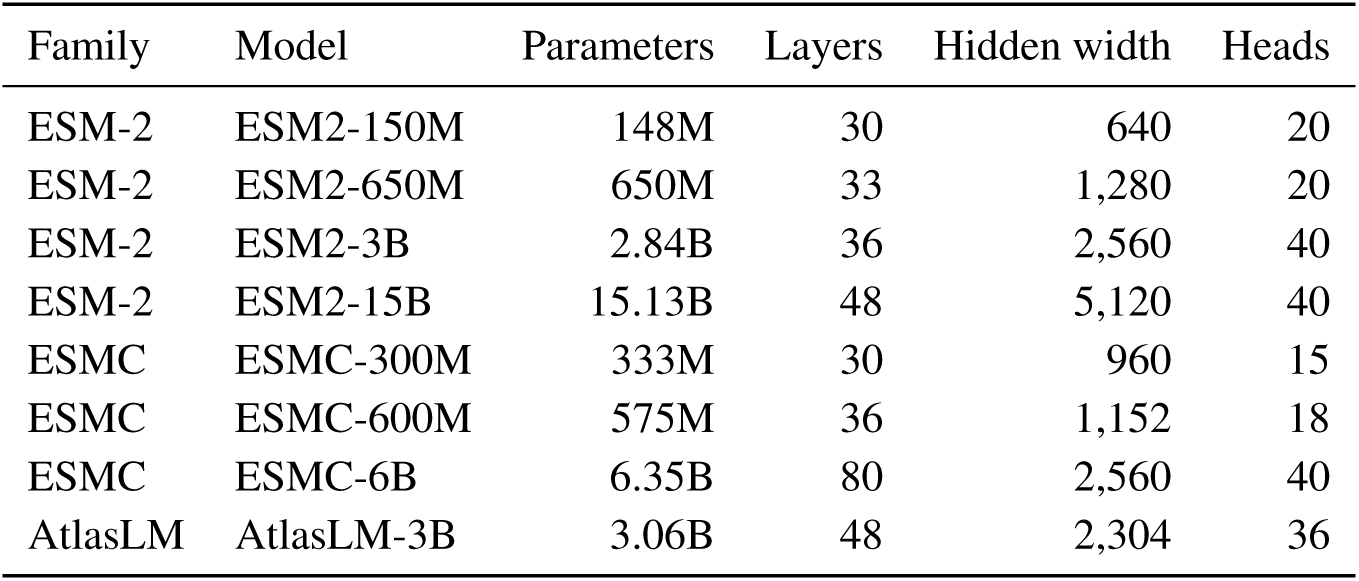
Model sizes included in the unsupervised contact experiment.

| Family | Model | Parameters | Layers | Hidden width | Heads |
| --- | --- | --- | --- | --- | --- |
| ESM-2 | ESM2-150M | 148M | 30 | 640 | 20 |
| ESM-2 | ESM2-650M | 650M | 33 | 1,280 | 20 |
| ESM-2 | ESM2-3B | 2.84B | 36 | 2,560 | 40 |
| ESM-2 | ESM2-15B | 15.13B | 48 | 5,120 | 40 |
| ESMC | ESMC-300M | 333M | 30 | 960 | 15 |
| ESMC | ESMC-600M | 575M | 36 | 1,152 | 18 |
| ESMC | ESMC-6B | 6.35B | 80 | 2,560 | 40 |
| AtlasLM | AtlasLM-3B | 3.06B | 48 | 2,304 | 36 |

#### A.2 Training

##### A.2.1 Training data

AtlasLM is trained on sequences prepared from UniRef (Suzek et al., 2007), MGnify (Mitchell et al., 2020), and MetaClust (Steinegger and Söding, 2018). The sequence-clustering and sampling procedure follows ESM3 (Hayes et al., 2025). Representative sequences are selected from clusters defined at 90% sequence identity, and training examples are sampled through collections clustered at 70% identity. Table 3 reports the prepared sequence and cluster counts used for sampling.

**Table 3:** AtlasLM training collections and 70%-identity cluster counts.

| Source | Sequences | Clusters |
| --- | --- | --- |
| UniRef | 188M | 83M |
| MGnify | 727M | 375M |
| MetaClust | 644M | 400M |

##### A.2.2 Optimization and training schedule

AtlasLM training uses masked-language modeling and AdamW with (*β*_1_*, β*_2_*, ɛ*) = (0.9, 0.95, 10*^−^*^8^), weight decay 0.01, peak learning rate 4 10*^−^*^4^, and gradient-norm clipping at 1.0. The first phase runs for 1.5M updates at context length 512 and global batch size 8,192, samples UniRef, MGnify, and MetaClust with proportions 36%, 32%, and 32%, and warms the learning rate for 5,000 updates. The second phase runs for 0.5M updates at context length 2,048 and global batch size 4,096, changes the sampling proportions to 62.5%, 25%, and 12.5%, and linearly decays the learning rate to 4 × 10*^−^*^5^.

##### A.2.3 Implementation and attention kernel

AtlasLM is implemented and trained with native PyTorch operators using PyTorch 2.10 or later. Variable-length sequences are packed and processed with the PyTorch `varlen_attn()` operator (PyTorch Foundation, 2026).

#### A.3 Unsupervised contact prediction protocol

The contact experiment evaluates language-model attention maps on partition 4 of the ESM structural superfamily split following the attention-probe protocol of Rao et al. (2021). Sequences longer than 510 residues are truncated to their first 510 residues. For every layer and attention head, special-token rows and columns are removed, the attention matrix is symmetrized, and average product correction is applied. Residue pairs separated by at least six positions are used to fit the probe. A pair is labeled as a structural contact when its reference distance is below 8 Å.

The probe is an L1-regularized logistic regression model implemented in scikit-learn, using *C* = 0.15, the `liblinear` solver, and at most 50 iterations. Each of ten independent repetitions samples 20 proteins without replacement from the 12,312-record training partition and fits an independent probe. At validation time, only long-range residue pairs satisfying *i j* 24 are ranked. The validation partition contains 2,985 records, of which 2,370 satisfy the implementation’s requirement of at least *L* evaluable long-range contacts and are scored in every repetition; the remaining 615 are skipped. P@L is the fraction of contacts among the top *L* pairs, and P@L/5 uses the top max(1*, L/*5) pairs. The values reported in the main-text figure are averages across the ten fitted probes.

Across AtlasLM-3B training, P@L increases from 0.5549 after 0.419 trillion processed tokens to 0.7147 after 10.486 trillion processed tokens (Fig. 9). This trajectory follows a single AtlasLM-3B training run; model-size scaling was not measured.

### B AtlasFold architecture, training, and evaluation

#### B.1 Architecture

##### B.1.1 Overview

AtlasFold predicts all-atom protein structures from a single amino-acid sequence without MSA or template inputs. It combines a frozen AtlasLM-3B encoder with a residue-level folding trunk and atom diffusion head. For a protein of length *L*, it maintains a single representation **s** R*^L×^*^384^ and pair representation **z** R*^L×L×^*^128^. Each recycling pass initializes these states from amino-acid identity and relative sequence position, adds recycled and LM features, and updates them with four LMStack blocks and a 48-block Pairformer. To reduce the parameter count, only the final 12 Pairformer blocks apply the pair-to-single update, comprising pair-biased self-attention and a single transition. The diffusion head is likewise specialized for protein atom14 coordinates, using reduced atom channels, attention heads, and residue-transformer depth. The final representations condition a 64-bin distogram head, diffusion sampling, and per-sample confidence heads. Algorithm 1 summarizes inference.

###### Algorithm 1

**AtlasFold inference**

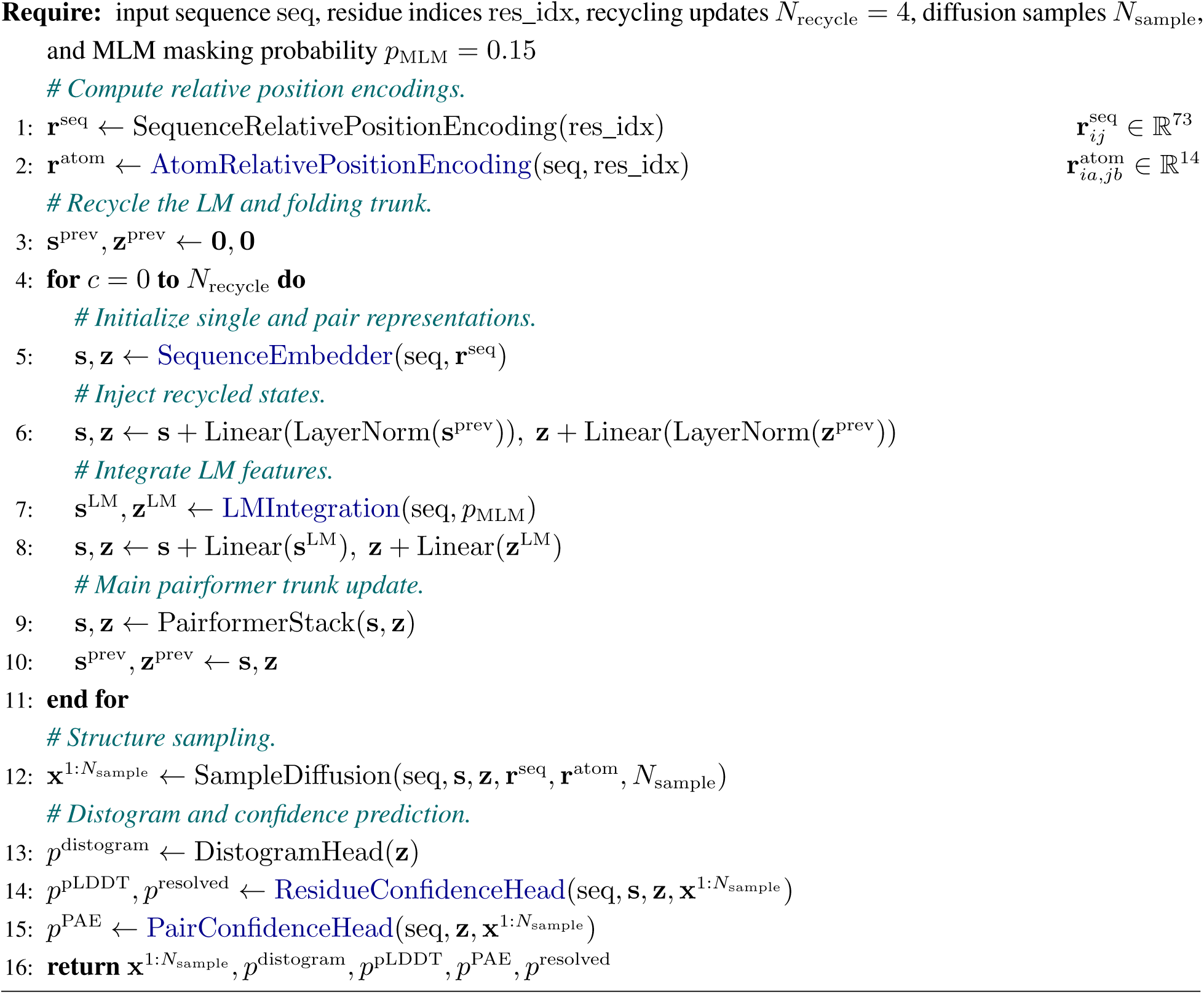

###### Algorithm 2

**Sequence embedding**

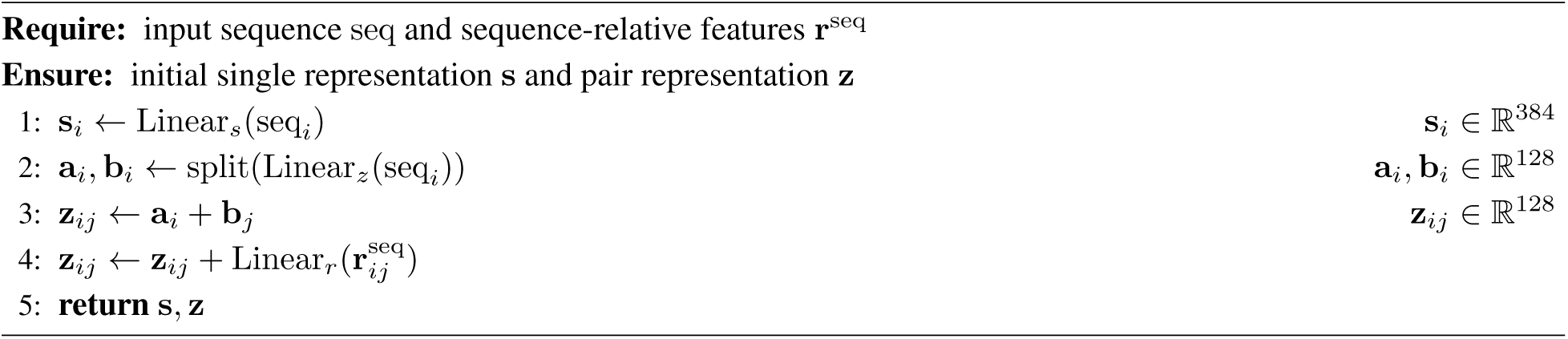

##### B.1.2 LM feature integration

The frozen LM returns hidden states and pre-softmax attention logits *A*^(*ℓ*)^ = *Q*^(*ℓ*)^(*K*^(*ℓ*)^)^T^*/*√*d_h_* from every transformer layer. The layer outputs are concatenated before a learned weighted sum forms the LM single feature and a learned projection forms the pair feature. The concatenation in Algorithm 3 is conceptual: for memory efficiency, the implementation projects each layer’s 36-head attention logits to 128 pair channels before accumulation, avoiding materialization of the concatenated attention tensor. After residue-token selection, separate MLPs produce a 768-channel LM single feature and a 128-channel pair feature. Four LMStack blocks then exchange information between these states through ESMFold-style single-to-pair, pair-to-pair, and pair-to-single updates (Lin et al., 2023).

###### Algorithm 3

**LM integration**

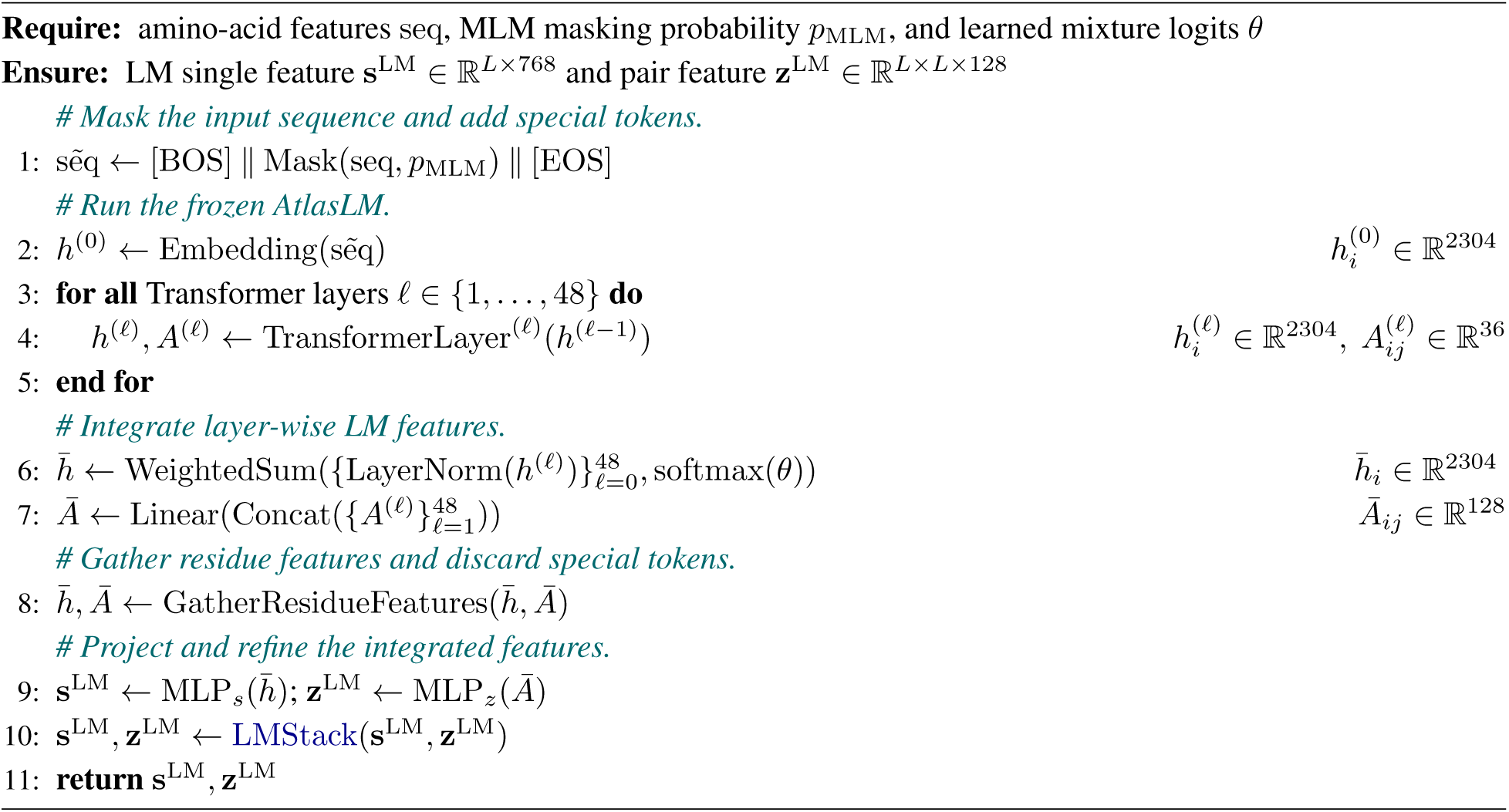

###### Algorithm 4

**LMStack**

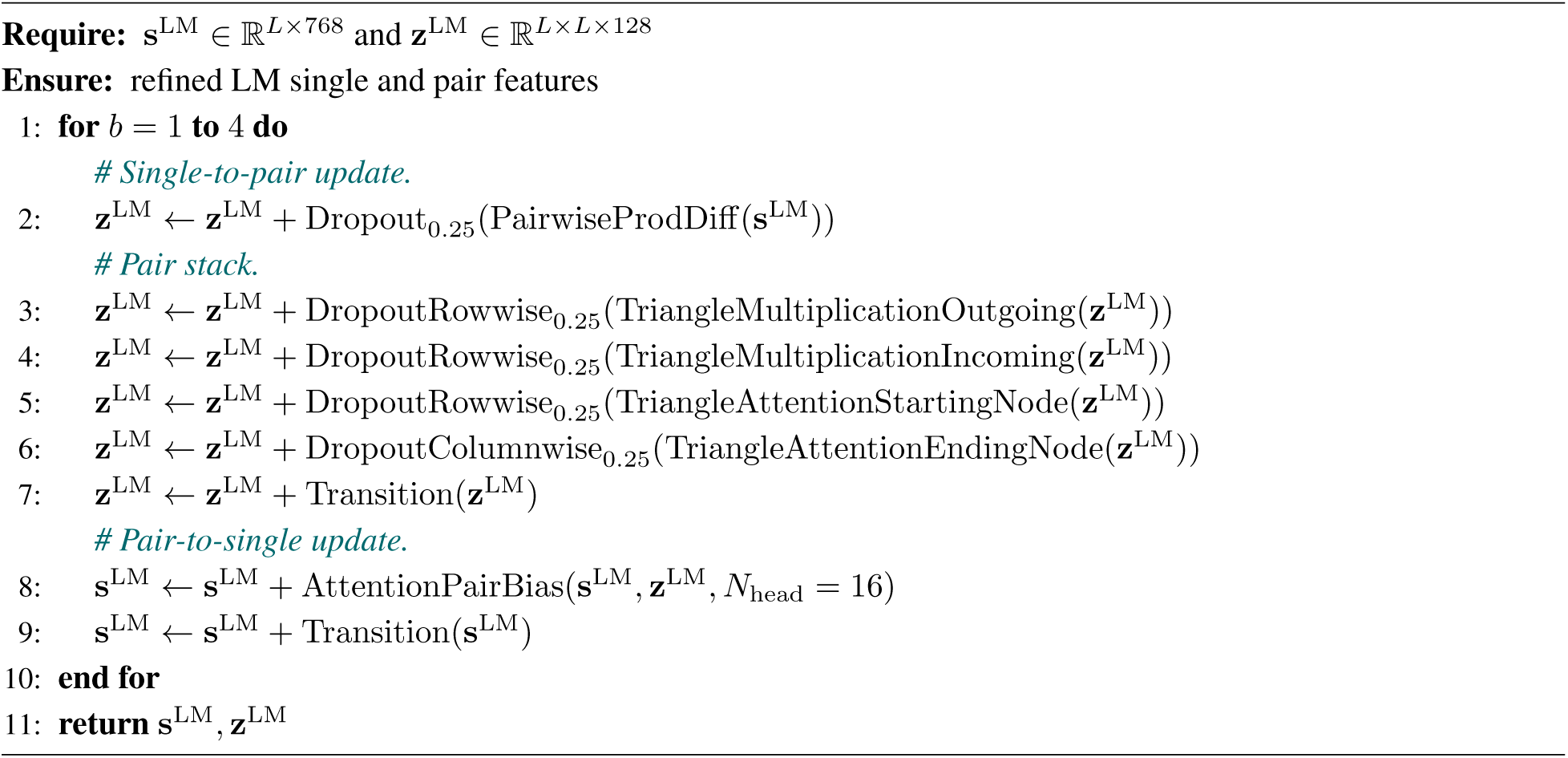

##### B.1.3 Atom-relative position encoding and diffusion head

The diffusion head uses EDM preconditioning and sampling (Karras et al., 2022). During sampling, every denoising step applies random centering, rotation, and translation to the coordinates. Its score model derives single conditioning from the trunk state, residue identity, and noise scale, and a global pair bias from the final pair representation and sequence-relative encoding (algorithm 6).

AlphaFold3 constructs an atom-pair representation *p_lm_* from per-atom metadata, trunk pair features, and geometry derived from a reference conformer, which is typically generated with RDKit ETKDGv3 and randomly rigidly augmented. Within its Atom Transformer, *p_lm_* is projected into the attention bias. By restricting atom modeling to proteins, AtlasFold instead uses a uniform residue-wise atom parameterization and replaces *p_lm_* with a deterministic **r**^atom^ that combines 10-channel clipped residue offsets with four-channel canonical intra-residue geometry before direct projection into block- and head-specific attention biases (algorithm 5).

This protein-specific design narrows atom features from 128 to 96 channels and local attention from four to two heads, and halves the global diffusion transformer from 24 to 12 blocks. Unlike AlphaFold3’s atom-index windows, AtlasFold masks attention by residue distance, retaining only same-chain atom pairs within ±4 residue positions (Abramson et al., 2024) (algorithms 7 and 8).

###### Algorithm 5

**Atom-relative position encoding**

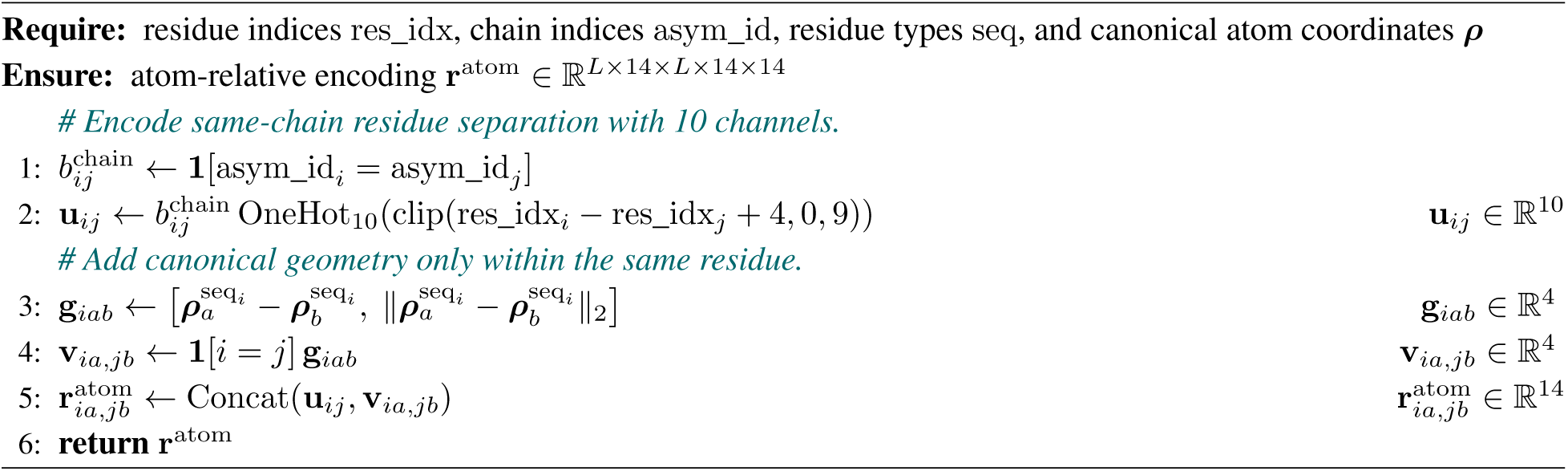

###### Algorithm 6

**Diffusion score model**

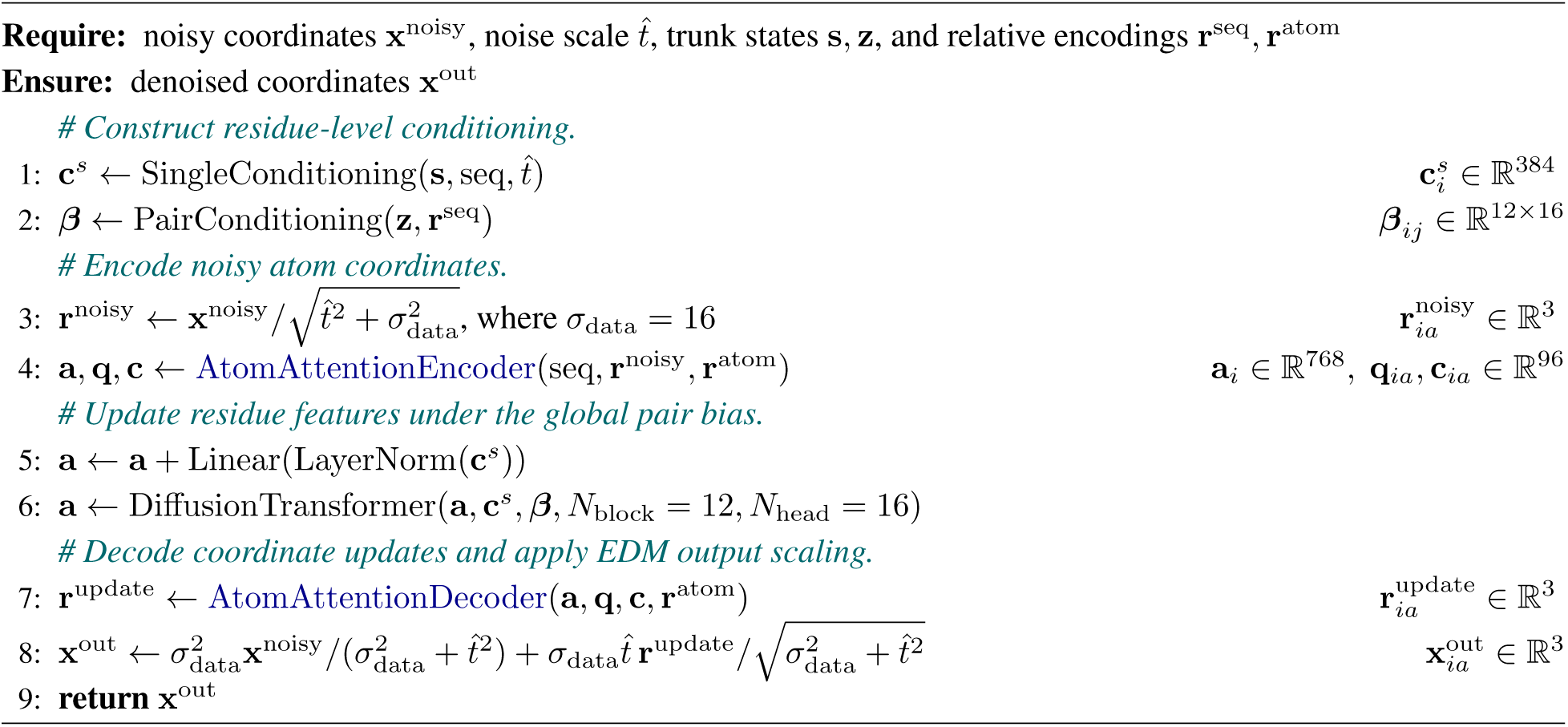

###### Algorithm 7

**Atom attention encoder**

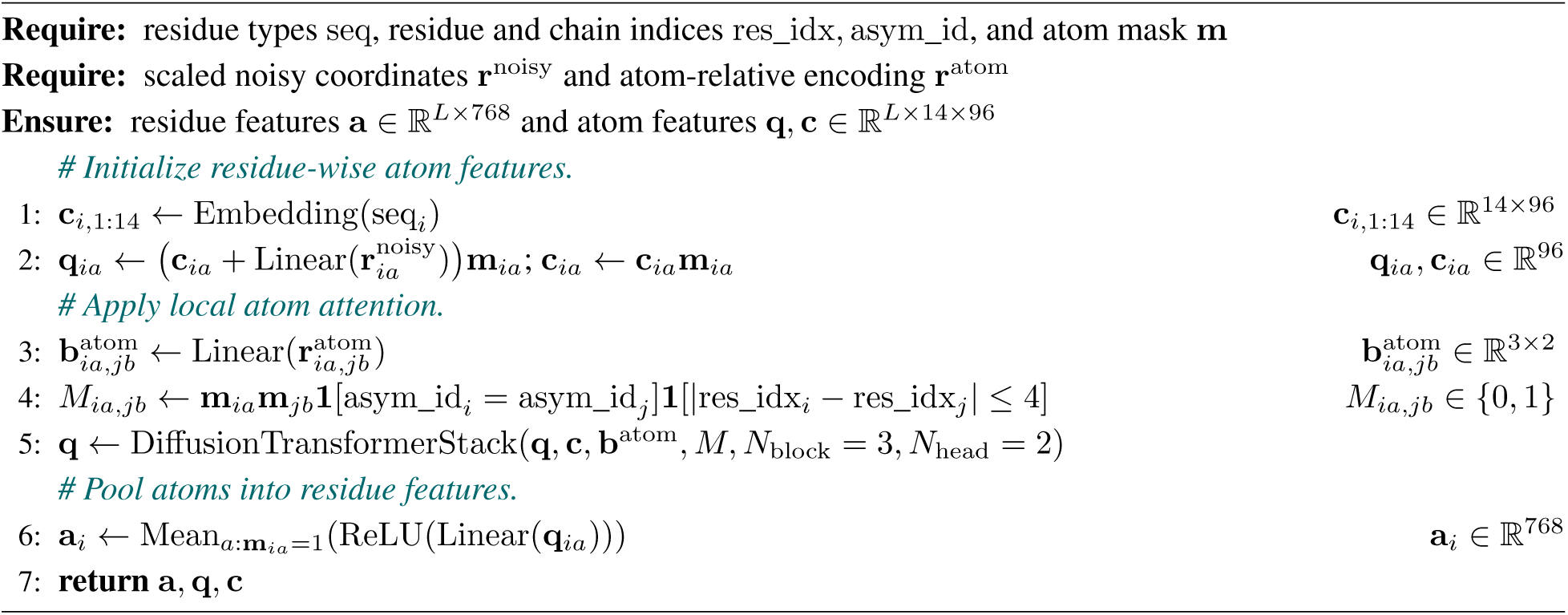

###### Algorithm 8

**Atom attention decoder**

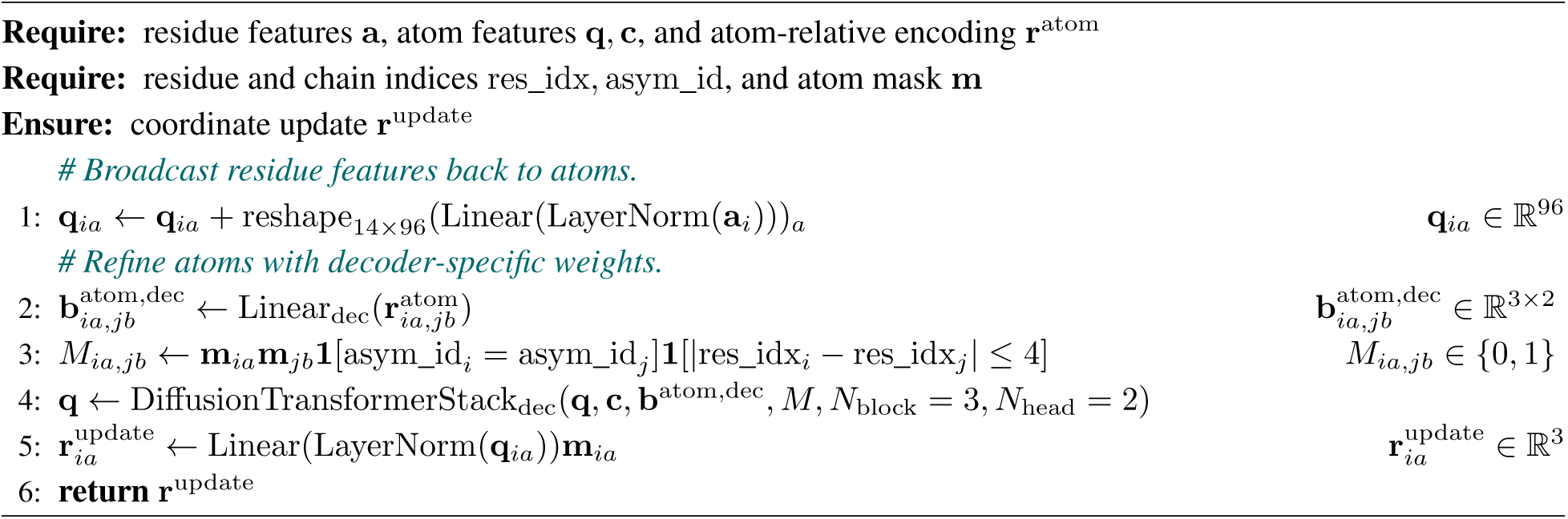

##### B.1.4 Confidence heads

For each sampled structure, both confidence heads detach the trunk states and coordinates, then augment the pair representation with residue-identity embeddings and a 39-bin pseudo-*C_β_* distogram computed from the sampled coordinates. The residue confidence head uses a two-block Pairformer to update the single representation and predicts pLDDT and experimentally resolved-atom probabilities (algorithm 9). The pair confidence head independently refines the augmented pair representation with a two-block pair stack and predicts PAE (algorithm 10).

###### Algorithm 9

**Residue confidence head**

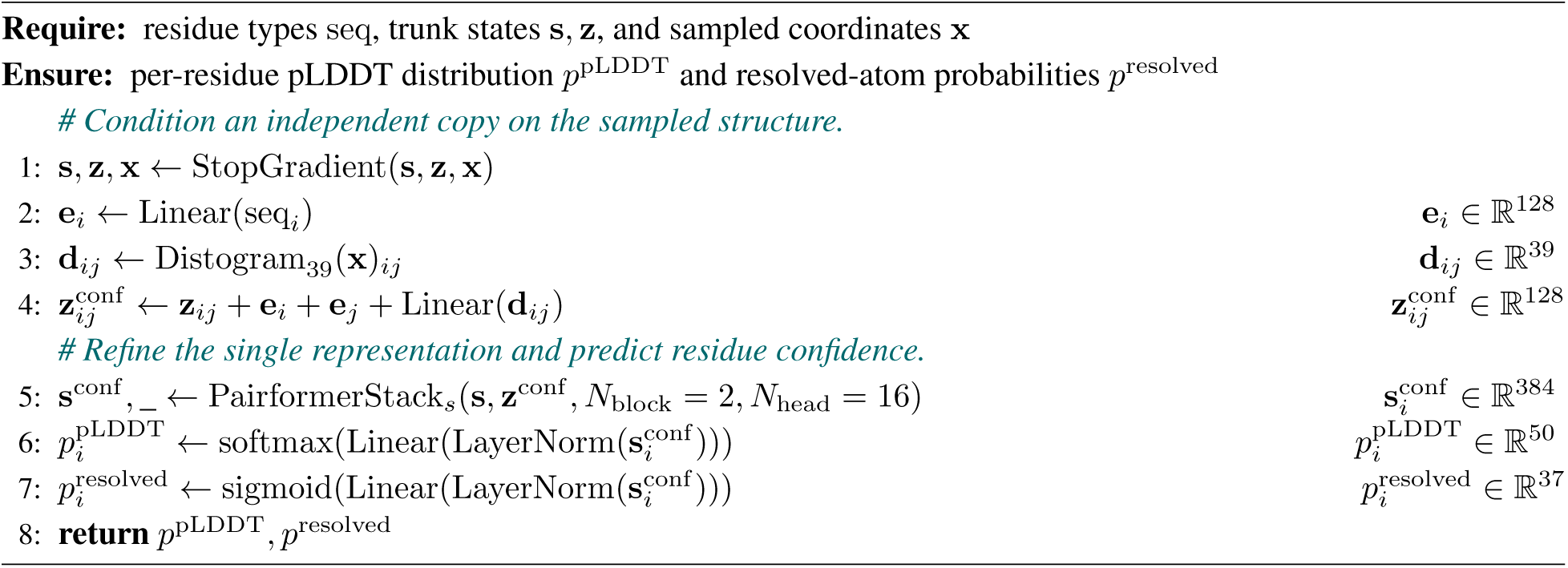

###### Algorithm 10

**Pair confidence head**

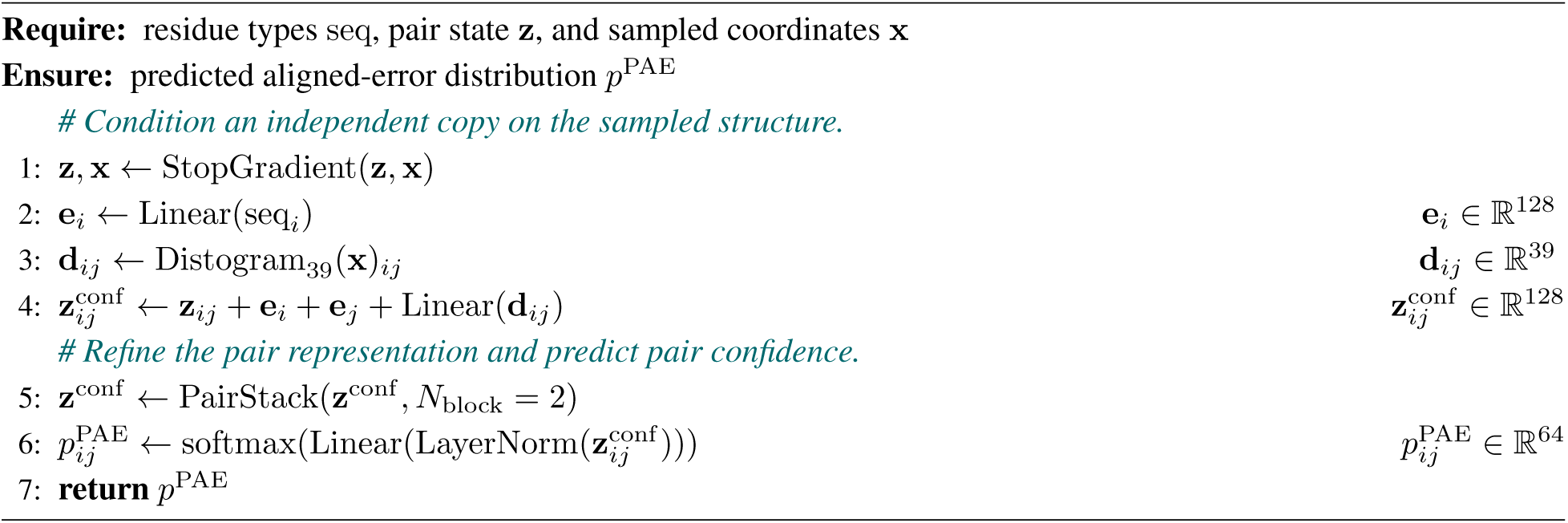

#### B.2 Training

##### B.2.1 Training data and validation set

Structural training draws from three sources:

1. **Experimental PDB structures.** The experimental training set contains Protein Data Bank (PDB) structures (Berman et al., 2000) released by 2020-05-01.
2. **MGnify-AF2.** The MGnify distillation structures are AlphaFold2 predictions distributed through the OpenFold Portal (Ahdritz et al., 2024).^3^ The source collection contains approximately 430,000 short sequences (fewer than 200 residues) and 16 million long sequences (at least 200 residues). After filtering predictions with mean pLDDT above 70, approximately 360,000 short and 15.4 million long sequences remain.
3. **Disordered PDB.** This source contains predicted structures for PDB chains with at least 40 unresolved residues. Stages 1 and 2 use five ESMFold predictions per sequence generated with a 15% input-masking ratio, retaining predictions with GDT-TS at least 60 over the experimentally resolved residues. After stage 2, we observed that these ESMFold predictions generally adopted packed conformations. This early exposure may contribute to the packing observed in IDR predictions from the current AtlasFold models. For stages 3 and 4, we therefore replaced them with predictions from AlphaFold2 pTM model 1. These predictions accounted for 1% of the training mixture in both stages, shifting the disordered-structure distribution toward AlphaFold2 structures. For new training runs, we recommend using the AlphaFold2 predictions from the outset rather than reproducing this intermediate dataset replacement; a smaller mixture proportion may be sufficient in this setting.

All training stages use the same temporally held-out CAMEO validation set used during ESMFold development (Lin et al., 2023). This validation set is used during model development and is distinct from the CAMEO22 benchmark and the confidence-evaluation set described in section 2.2.2.

##### B.2.2 Dataset sampling

MMseqs2 clusters are constructed at 40% sequence identity. Cluster weighting is inversely proportional to cluster size. Following AlphaFold2 (Jumper et al., 2021), length weighting uses clip(*L,* 256, 512). Distillation confidence weighting uses clip(pLDDT − 30, 0, 40).

##### B.2.3 Cropping

When a protein exceeds the current crop length, AtlasFold selects a spatial crop with probability 0.6, a contiguous crop with probability 0.2, or a multi-contiguous crop with probability 0.2. Spatial cropping selects residues nearest to a valid *C_α_* anchor and falls back to contiguous cropping when geometry is unavailable. Multi-contiguous cropping draws two to four sequence segments, allocates at least 32 residues to each selected segment, samples one interval within each segment, and restores sequence order after concatenation. This procedure exposes separated regions of long proteins within a fixed crop length.

##### B.2.4 Training objectives

The monomer objective is

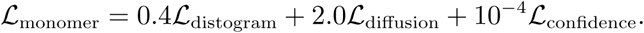

The diffusion term is a noise-weighted coordinate mean-squared error. During stages in which the smooth-lDDT option is enabled, its loss is added once:

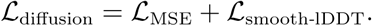

The smooth-lDDT configuration value acts as an enable gate and is not applied as a second multiplier. The monomer confidence loss is

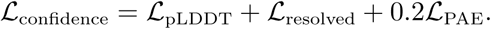

Confidence losses are enabled only for non-distilled X-ray or cryo-EM structures with recorded resolution from 0.1 to 3.0 Å.

##### B.2.5 Optimization and training schedule

Structure training uses Adam with (*β*_1_*, β*_2_*, ɛ*) = (0.9, 0.95, 10*^−^*^8^), mixed BF16 precision, global batch size 256, diffusion batch size 32, up to three recycling updates sampled per training step, exponential moving average decay 0.999, and gradient-norm clipping at 5.0. For the first three stages, the learning rate warms up for 1,000 updates to a maximum of 1.8 10*^−^*^3^ and decays by a factor of 0.95 every 50,000 updates. AtlasLM is excluded from optimization and from the exponential moving average. The crop and sequence limits increase across stages (Table 4). The first two stages use the smooth-lDDT term. The final 2k-update stage freezes the trunk and uses a fixed learning rate of 10*^−^*^3^. Table 4 reproduces the checkpoint-to-checkpoint dataset mixtures used during training.

###### Confidence-head retraining

With an AlphaFold3-style shared confidence module, we observed that pLDDT-based sample ranking degraded after PAE supervision was introduced during fine-tuning. Therefore, we split the module into separate local and pairwise confidence paths and retrained the confidence module from scratch after stage 4. For new training runs, this retraining procedure can be omitted by using the separated confidence architecture from the start of training.

**Table 4:** AtlasFold monomer training stages.

| Setting | Stage 1 | Stage 2 | Stage 3 | Stage 4 |
| --- | --- | --- | --- | --- |
| Updates | 45k | 45k | 8k | 2k |
| Crop length | 256 | 384 | 512 | 640 |
| Sequence limit | 512 | 768 | 1,024 | 1,280 |
| RCSB PDB (%) | 12.0 | 12.3 | 24.0 | 24.0 |
| Disordered PDB (%) | 0.5 | 0.2 | 1.0 | 1.0 |
| MGNify-AF2-long (%) | 86.5 | 86.5 | 74.0 | 74.0 |
| MGNify-AF2-short (%) | 1.0 | 1.0 | 1.0 | 1.0 |
| Length-based sampling | O | O | X | X |
| Smooth-IDDT loss | O | O | X | X |
| Train trunk | O | O | O | X |

#### B.3 Evaluation protocol

The benchmark sets contain 183 CAMEO22, 70 CASP14, and 56 CASP15 proteins. TM-score (Zhang and Skolnick, 2004), GDT-TS, lDDT (Mariani et al., 2013), lDDT-*C_α_*, and RMSD are computed with OpenStructure 2.9.1 (Biasini et al., 2013).

##### B.3.1 Baselines and inference settings

###### AlphaFold2

AlphaFold2 (Jumper et al., 2021) uses ColabFold (Mirdita et al., 2022) with MMseqs2 (Steinegger and Söding, 2017) MSAs, three recycles, and no templates or Amber relaxation.

###### RoseTTAFold

RoseTTAFold results are reproduced from SimpleFold (Wang et al., 2026).

###### RoseTTAFold2

RoseTTAFold2 (Baek et al., 2023) uses ColabFold with 3 recycles, 256 MSA sequences, and no subcropping.

###### ESMFold

ESMFold (Lin et al., 2023) uses the `esmfold_v1` checkpoint with 4 recycles. An optimized implementation^4^ with cuEquivariance kernels (NVIDIA, 2026) is used.

###### SimpleFold-3B

SimpleFold-3B (Wang et al., 2026) uses the released 3B-parameter checkpoint with *τ* = 0.01 and 500 diffusion steps.

###### ESMFold2

ESMFold2 (Candido et al., 2026) uses PyTorch 2.9.1, NVIDIA Transformer Engine 2.13.0, and cuEquivariance 0.10.0. It uses FP8 for ESMC. Default inference uses 10 loops, 5 diffusion samples, and 68-step truncated diffusion without an MSA.

###### AtlasFold

AtlasFold uses PyTorch 2.10.0 and cuEquivariance 0.10.0. Default inference uses 4 recycles, 5 diffusion samples, and a length-adaptive schedule of 20, 30, or 100 steps for lengths up to 512, 1,024, or above 1,024, respectively.

Table 5 summarizes candidate generation and model selection for the monomer aggregate comparison.

**Table 5:** Prediction generation and selection used for the monomer aggregate table.

| Model | Candidate generation | Selection |
| --- | --- | --- |
| RoseTTAFold | single prediction | – |
| RoseTTAFold2 | single prediction | – |
| ESMFold | single prediction | – |
| AlphaFold2 | five model outputs | average pLDDT |
| SimpleFold-3B | five diffusion samples | average pLDDT |
| ESMFold2 | seeds 1–5, one sample per seed | average pLDDT |
| AtlasFold | seeds 1–5, one sample per seed | average pLDDT |

**Table 6:** PDB structural-training cutoff dates used to interpret the monomer benchmarks.

| Model | PDB cutoff | Model | PDB cutoff |
| --- | --- | --- | --- |
| AlphaFold2 | 2018-05-01 | ESMFold | 2020-05-01 |
| RoseTTAFold | 2020-02-17 | SimpleFold | 2020-05-01 |
| RoseTTAFold2 | 2020-04-30 | AtlasFold | 2020-05-01 |
| ESMFold2 | 2021-09-30 |  |  |

<sup>4</sup>https://github.com/SeonghwanSeo/esmfold-minimal

### C AtlasFold-Multimer architecture, fine-tuning, and evaluation

#### C.1 Architecture changes

AtlasFold-M is fine-tuned from AtlasFold. The language-model, initialization, Pairformer, distogram, and diffusion modules are retained, while the template module and AlphaFold3-style confidence module are introduced for complex training. The template module contains two blocks with channel width 64 and a 39-bin template distogram. The confidence module contains four blocks and predicts pLDDT, PAE, PDE, and resolved distributions and derives pTM, ipTM, chain pTM, and pairwise interface ipTM.

Before entering the multimer trunk, AtlasLM attention-derived pair features are masked to retain only intra-chain residue pairs. Inter-chain pair features are then formed and refined by the subsequent triangular update modules. Final structures are ranked by 0.8 ipTM + 0.2 pTM + 0.5 disorder, following the AlphaFold3 confidence-ranking notation (Abramson et al., 2024). Here, disorder is the fraction of residues classified as disordered from the predicted relative solvent-accessible surface area.

**Table 7:** AtlasFold-M architecture and outputs that differ from, or extend, AtlasFold monomer.

| Component | Configuration |
| --- | --- |
| Template channel / blocks | 64 / 2 |
| Template distogram bins | 39 |
| Maximum template slots per chain | 2 |
| Confidence blocks | 4 |
| pLDDT / PAE / PDE bins | 50 / 64 / 64 |
| Ranking score | $0.8 \text{ ipTM} + 0.2 \text{ pTM} + 0.5 \text{ disorder}$ |

#### C.2 Fine-tuning

##### C.2.1 Training data and validation set

Multimer fine-tuning draws from three sources:

1. **Experimental PDB complexes.** The experimental training set contains PDB complexes (Berman et al., 2000) released by 2021-09-30. Preprocessing expands biological assemblies, validates geometry, filters all-atom clashes, and identifies interfaces. Assemblies with more than 20 valid protein chains are reduced to a 20-chain subcomplex around a sampled interface or contact seed.
2. **Disordered PDB.** This source contains AlphaFold-Multimer predictions for PDB sequences containing disordered regions.
3. **MGnify-AF2.** The monomer MGnify distillation sets described in Section B.2.1 are retained during multimer fine-tuning.

Following AlphaFold3 (Abramson et al., 2024), the multimer validation set is drawn from PDB structures released after 2021-09-30 and before 2023-01-13 and selected through low-homology interface filtering. AtlasFold-M retains protein complexes with 2–20 chains, 16–1,536 protein residues, and at least one interface, rather than AlphaFold3’s 2,048-token limit. An interface is retained when no training complex contains homologs at 40% sequence identity to both partners; the retained interfaces are clustered by their two chain-cluster identifiers, one interface is sampled per cluster, and the resulting targets are subsampled to 512 complexes.

##### C.2.2 Dataset sampling and templates

Protein chains are clustered with MMseqs2 at 40% sequence identity, and an interface cluster is identified by the pair of its two chain-cluster identifiers. Within the experimental-complex dataset, either a chain item or an interface item is sampled with weight *w*_type_*/N*_cluster_, where *N*_cluster_ is the number of items sharing its cluster identifier, *w*_chain_ = 1, and *w*_interface_ = 2.

Only the experimental-complex source uses templates during training. Template hits must have been released at least 60 days before the target entry. With probability 0.4, one or two available hits are sampled uniformly up to the two-slot limit; otherwise, both slots are empty. Template selection is performed per query chain, and aligned atom14 template features are concatenated across chains. The template database was constructed from template resources downloaded from the OpenFold Portal (Ahdritz et al., 2024).^5^ Validation and the reported evaluation on the two FoldBench subsets are template free.

##### C.2.3 Cropping

Experimental multimer examples use spatial, interface-spatial, and contiguous crops with probabilities 0.4, 0.4, and 0.2. Interface-spatial cropping selects an interface residue as the geometric anchor and falls back to spatial cropping when no valid interface anchor exists. Contiguous cropping first samples a chain, orders nearby chains by spatial proximity, and considers at most the stage-specific maximum number of chains. The maximum is six chains in stages 1 and 2 and eight chains in stage 3. Monomer distillation examples included during fine-tuning use spatial and contiguous crops with probabilities 0.75 and 0.25.

##### C.2.4 Training objectives

AtlasFold-M uses the same loss components and global weights as AlphaFold3 (Abramson et al., 2024):

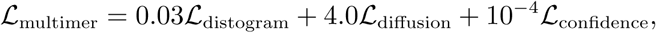

where the stage-1 diffusion term includes smooth-lDDT and the multimer confidence term is

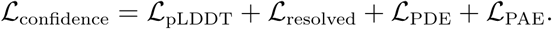

Unlike AlphaFold3, which activates the PAE term only in its final training stage, AtlasFold-M applies the PAE term throughout all three fine-tuning stages. As for monomer training, confidence losses are applied only to non-distilled X-ray or cryo-EM structures with recorded resolution from 0.1 to 3.0 Å. Before computing these losses, ground-truth chains and symmetry-equivalent assignments are aligned to the confidence mini-rollout prediction.

##### C.2.5 Optimization and training schedule

Fine-tuning uses Adam with the AtlasFold optimizer parameters, mixed BF16 precision, global batch size 256, diffusion batch size 32, up to three recycling updates sampled per training step, exponential moving average decay 0.999, and gradient-norm clipping at 5.0. AtlasFold-M is trained for 25,000 additional updates in three stages (Table 8). For the first two stages, the learning rate warms up for 1,000 updates to a maximum of 1.8 10*^−^*^3^ and decays by a factor of 0.95 every 50,000 updates. The final stage freezes the trunk, initializes it from exponential-moving-average weights, and uses a fixed learning rate of 10*^−^*^3^.

**Table 8:** AtlasFold-M fine-tuning stages.

| Setting | Stage 1 | Stage 2 | Stage 3 |
| --- | --- | --- | --- |
| Updates | 21k | 2k | 2k |
| Crop length | 384 | 640 | 768 |
| Sequence limit | 768 | 1,280 | 1,536 |
| Maximum chains | 6 | 6 | 8 |
| RCSB PDB (%) | 73.0 | 49.0 | 49.0 |
| Disordered PDB (%) | 2.0 | 1.0 | 1.0 |
| Mgnify-AF2-long (%) | 24.5 | 49.5 | 49.5 |
| Mgnify-AF2-short (%) | 0.5 | 0.5 | 0.5 |
| Smooth-IDDT loss | O | X | X |
| Train trunk | O | O | X |

#### C.3 Evaluation protocol

##### C.3.1 Baselines and inference settings

AlphaFold3, Boltz-1, Protenix-v1, and ESMFold2-MSA use MSAs prepared through the AlphaFold3 input pipeline. ESMFold2 and AtlasFold-M are evaluated without MSAs.

###### AlphaFold-Multimer v2.3

AlphaFold-Multimer v2.3 (Evans et al., 2021) is run with ColabFold default settings, `num_recycles=10`, and `tol=0`.

###### AlphaFold3

AlphaFold3 (Abramson et al., 2024) version 3.0.1 uses 10 recycles and 200 diffusion steps.

###### Boltz-1

Boltz-1 (Wohlwend et al., 2024) uses 10 recycles and 200 diffusion steps. Boltz-1 is run from commit `b1ebfc4` with PyTorch 2.10.0 and cuEquivariance 0.10.0.

###### Protenix-v1

Protenix-v1 (Zhang et al., 2026) uses 10 recycles and 200 diffusion steps. Protenix-v1 is run from commit `4c355be` with PyTorch 2.10.0 and cuEquivariance 0.8.0.

###### ESMFold2 and ESMFold2-MSA

ESMFold2 and ESMFold2-MSA use PyTorch 2.9.1, NVIDIA Transformer Engine 2.13.0, and cuEquivariance 0.10.0. Both modes use FP8 for ESMC and run 10 loops with 68-step truncated diffusion.

###### AtlasFold-M

AtlasFold-M uses PyTorch 2.10.0 and cuEquivariance 0.10.0. It uses 10 recycles, an MLM masking probability of 0.20, 200 diffusion steps, and no templates.

##### C.3.2 FoldBench benchmark results

The interface evaluation covers all 172 Ab–Ag targets and 278 of the 279 P–P targets from FoldBench (Xu et al., 2025); the excluded P–P target, 8GQP, is a D-protein binder system. Model-specific FoldBench settings are summarized in Section C.3.1. For the diffusion-based methods, we generate five diffusion samples per seed, whereas AlphaFold-Multimer v2.3 generates one prediction from each of five models per seed. To estimate performance with five seeds while reducing sensitivity to the particular seeds selected, we generate predictions with seeds 1–10 for all seven methods and evaluate every five-seed subset. This yields ^10^ = 252 combinations of 25 candidates.

Figure 5 reports Top-1, Oracle, and Avg. summaries. Top-1 selects a structure using each model’s native confidence-ranking score, Oracle selects the generated structure with the highest DockQ against the reference structure, and Avg. averages across all generated structures. For each summary and metric, we report the mean across the 252 seed combinations; the error bars in Fig. 5 denote one standard deviation of the acceptable-or-better success rate across these combinations. DockQ scores (Basu and Wallner, 2016) are computed with OpenStructure 2.9.1 (Biasini et al., 2013), using acceptable, medium, and high thresholds of 0.23, 0.49, and 0.80. Oracle is unavailable at inference time because it requires the reference structure. For AlphaFold-Multimer v2.3 on P–P, DockQ-derived metrics, Fnat, iRMSD, and LRMSD are available for 278 of 279 targets, whereas lDDT is available for all 279 targets. Tables 9 and 10 report the corresponding complete metric-set means from the evaluation output.

**Table 9:** FoldBench antibody–antigen results. High, Medium+, and Accept.+ report the fractions above DockQ thresholds of 0.80, 0.49, and 0.23, respectively; higher is better except for iRMSD and LRMSD.

| Selection | Model | High | Medium+ | Accept.+ | DockQ | Fnat | iRMSD | LRMSD | IDDT |
| --- | --- | --- | --- | --- | --- | --- | --- | --- | --- |
| Rank | AF-Multimer v2.3 | 8.516 | 27.935 | 40.924 | 0.276 | 0.275 | 8.221 | 27.611 | 0.797 |
|  | AlphaFold3 | 20.951 | 39.636 | 49.285 | 0.376 | 0.383 | 7.267 | 23.962 | 0.877 |
|  | Boltz-1 | 5.987 | 22.566 | 34.967 | 0.235 | 0.240 | 8.734 | 28.759 | 0.841 |
|  | Protenix-v1 | 21.288 | 37.394 | 44.945 | 0.354 | 0.362 | 7.477 | 23.275 | 0.866 |
|  | ESMFold2-MSA | 21.036 | 42.693 | 52.270 | 0.391 | 0.401 | 7.165 | 24.615 | 0.869 |
|  | ESMFold2 | 20.981 | 42.290 | 48.602 | 0.378 | 0.394 | 7.690 | 25.724 | 0.849 |
|  | AtlasFold-M | 18.173 | 35.359 | 46.955 | 0.346 | 0.361 | 7.616 | 25.631 | 0.861 |
| Oracle | AF-Multimer v2.3 | 11.213 | 34.542 | 52.921 | 0.355 | 0.395 | 5.365 | 17.155 | 0.826 |
|  | AlphaFold3 | 29.614 | 49.285 | 66.168 | 0.486 | 0.519 | 4.009 | 12.203 | 0.891 |
|  | Boltz-1 | 10.341 | 24.958 | 39.505 | 0.283 | 0.310 | 6.870 | 21.137 | 0.853 |
|  | Protenix-v1 | 28.899 | 46.099 | 65.818 | 0.467 | 0.505 | 4.247 | 12.771 | 0.882 |
|  | ESMFold2-MSA | 33.230 | 49.059 | 63.427 | 0.480 | 0.513 | 4.100 | 11.747 | 0.881 |
|  | ESMFold2 | 31.467 | 48.491 | 61.489 | 0.474 | 0.516 | 4.196 | 11.717 | 0.862 |
|  | AtlasFold-M | 22.626 | 47.628 | 64.733 | 0.460 | 0.512 | 4.142 | 13.203 | 0.874 |
| Sample avg. | AF-Multimer v2.3 | 3.349 | 12.628 | 24.477 | 0.175 | 0.169 | 9.689 | 31.560 | 0.772 |
|  | AlphaFold3 | 17.988 | 35.523 | 43.547 | 0.339 | 0.344 | 7.686 | 25.063 | 0.875 |
|  | Boltz-1 | 5.663 | 19.953 | 32.221 | 0.220 | 0.227 | 9.021 | 29.969 | 0.840 |
|  | Protenix-v1 | 16.802 | 32.035 | 40.779 | 0.311 | 0.320 | 8.083 | 26.182 | 0.863 |
|  | ESMFold2-MSA | 18.733 | 36.837 | 44.709 | 0.342 | 0.350 | 7.949 | 26.945 | 0.862 |
|  | ESMFold2 | 16.930 | 34.802 | 42.395 | 0.323 | 0.336 | 8.847 | 29.476 | 0.840 |
|  | AtlasFold-M | 13.267 | 27.640 | 38.453 | 0.283 | 0.284 | 8.585 | 28.197 | 0.855 |

**Table 10:** FoldBench protein–protein results for 278 targets. Metrics and selection modes follow Table 9.

| Selection | Model | High | Medium+ | Accept.+ | DockQ | Fnat | iRMSD | LRMSD | IDDT |
| --- | --- | --- | --- | --- | --- | --- | --- | --- | --- |
| Rank | AF-Multimer v2.3 | 37.416 | 59.244 | 65.206 | 0.515 | 0.542 | 8.144 | 17.005 | 0.718 |
|  | AlphaFold3 | 43.424 | 68.334 | 73.006 | 0.592 | 0.641 | 5.656 | 13.852 | 0.844 |
|  | Boltz-1 | 37.316 | 62.229 | 71.120 | 0.551 | 0.598 | 6.015 | 14.499 | 0.800 |
|  | Protenix-v1 | 44.043 | 66.966 | 74.445 | 0.588 | 0.637 | 5.585 | 13.790 | 0.832 |
|  | ESMFold2-MSA | 42.983 | 66.638 | 73.230 | 0.587 | 0.637 | 5.817 | 14.183 | 0.806 |
|  | ESMFold2 | 35.703 | 61.786 | 69.216 | 0.537 | 0.586 | 6.418 | 15.016 | 0.791 |
|  | AtlasFold-M | 26.829 | 54.291 | 63.358 | 0.474 | 0.517 | 7.570 | 18.198 | 0.768 |
| Oracle | AF-Multimer v2.3 | 43.294 | 63.312 | 68.615 | 0.565 | 0.611 | 6.252 | 12.649 | 0.757 |
|  | AlphaFold3 | 55.488 | 72.499 | 78.028 | 0.659 | 0.712 | 3.882 | 8.766 | 0.859 |
|  | Boltz-1 | 46.834 | 66.505 | 76.632 | 0.605 | 0.664 | 4.612 | 10.728 | 0.816 |
|  | Protenix-v1 | 54.832 | 74.499 | 81.572 | 0.666 | 0.723 | 3.674 | 8.284 | 0.852 |
|  | ESMFold2-MSA | 55.434 | 71.363 | 77.237 | 0.655 | 0.715 | 3.942 | 8.506 | 0.823 |
|  | ESMFold2 | 47.917 | 66.906 | 74.071 | 0.613 | 0.669 | 4.186 | 8.506 | 0.809 |
|  | AtlasFold-M | 40.636 | 64.314 | 71.560 | 0.576 | 0.635 | 4.669 | 10.627 | 0.799 |
| Sample avg. | AF-Multimer v2.3 | 32.065 | 54.281 | 59.698 | 0.471 | 0.498 | 8.649 | 18.553 | 0.693 |
|  | AlphaFold3 | 43.640 | 66.561 | 71.504 | 0.582 | 0.626 | 5.833 | 14.119 | 0.841 |
|  | Boltz-1 | 36.770 | 60.942 | 70.230 | 0.538 | 0.587 | 6.268 | 15.188 | 0.791 |
|  | Protenix-v1 | 40.986 | 60.719 | 69.144 | 0.545 | 0.584 | 6.359 | 15.625 | 0.815 |
|  | ESMFold2-MSA | 38.712 | 62.662 | 68.971 | 0.550 | 0.600 | 6.487 | 15.880 | 0.800 |
|  | ESMFold2 | 31.978 | 58.547 | 65.712 | 0.507 | 0.556 | 7.087 | 17.059 | 0.784 |
|  | AtlasFold-M | 26.971 | 52.496 | 60.676 | 0.456 | 0.496 | 7.965 | 19.632 | 0.766 |

### D Computational profiling

#### D.1 Inference settings

All computational profiles were collected on a single NVIDIA B200 GPU with CUDA 12.8, and reported latencies exclude MSA-search time. Model-specific software versions and inference settings are provided in Sections B.3.1 and C.3.1.

#### D.2 Monomer inference profiling

Table 11 compares peak GPU memory for AtlasFold, ESMFold2, ESMFold, SimpleFold-3B, and AlphaFold2 across sequence lengths. Table 12 reports monomer inference runtime across the same sequence lengths. Table 13 reports the five-sample settings plotted in Fig. 7a.

**Table 11:** Peak GPU memory during inference.

| Length | Peak GPU memory (GiB) |  |  |  |  |
| --- | --- | --- | --- | --- | --- |
|  | AtlasFold | ESMFold2 | ESMFold | SimpleFold-3B | AlphaFold2 |
| 64 | 6.74 | 7.12 | 7.91 | 21.48 | 3.81 |
| 128 | 6.82 | 7.54 | 7.98 | 21.56 | 4.06 |
| 192 | 6.99 | 8.21 | 8.11 | 21.65 | 4.19 |
| 256 | 7.22 | 9.26 | 8.29 | 21.74 | 4.43 |
| 384 | 7.88 | 12.26 | 8.79 | 21.96 | 4.81 |
| 512 | 8.80 | 16.46 | 9.49 | 22.23 | 5.31 |
| 640 | 9.97 | 21.86 | 10.39 | 22.56 | 6.19 |
| 768 | 11.40 | 28.46 | 11.48 | 22.88 | 6.94 |
| 896 | 13.08 | 36.25 | 12.78 | 23.29 | 7.56 |
| 1,024 | 15.02 | 45.24 | 14.27 | 23.70 | 8.32 |
| 1,280 | 19.68 | 66.82 | 17.84 | 24.63 | 9.07 |
| 1,536 | 25.36 | 93.19 | 22.21 | 25.88 | 9.94 |
| 1,792 | 32.07 | 124.36 | 27.37 | 27.42 | 12.94 |
| 2,048 | 39.81 | 160.32 | 33.32 | 29.44 | 16.57 |

**Table 12:** Monomer inference runtime without MSA search in seconds per sequence.

| $L$ | AtlasFold | | ESMFold2 | | SimpleFold-3B | ESMFold | AlphaFold2 |
| --- | --- | --- | --- | --- | --- | --- | --- |
|  | 1 sample | 5 samples | 1 sample | 5 samples | 1 sample | 1 model | 1 model |
| 64 | 1.18 | 1.29 | 1.99 | 2.21 | 27.04 | 0.77 | 3.10 |
| 128 | 1.38 | 1.50 | 2.53 | 2.75 | 28.80 | 0.98 | 3.96 |
| 192 | 1.39 | 1.49 | 2.65 | 3.18 | 29.98 | 0.98 | 5.04 |
| 256 | 1.39 | 1.50 | 2.78 | 3.76 | 34.97 | 1.08 | 5.86 |
| 384 | 1.57 | 1.74 | 3.73 | 6.24 | 40.73 | 2.23 | 9.86 |
| 512 | 2.41 | 2.76 | 5.77 | 9.89 | 50.00 | 4.02 | 12.44 |
| 640 | 3.76 | 4.49 | 8.46 | 14.78 | 60.11 | 6.82 | 16.85 |
| 768 | 5.35 | 6.33 | 11.85 | 19.94 | 68.43 | 10.33 | 21.23 |
| 896 | 7.41 | 8.70 | 17.65 | 27.90 | 80.68 | 15.23 | 27.49 |
| 1,024 | 9.90 | 11.57 | 25.53 | 39.17 | 88.01 | 21.04 | 34.38 |
| 1,280 | 18.25 | 24.42 | 43.71 | 61.93 | 111.99 | 36.80 | 51.85 |
| 1,536 | 27.53 | 35.82 | 67.00 | 92.67 | 140.69 | 58.53 | 74.36 |
| 1,792 | 40.14 | 50.28 | 99.02 | 132.85 | 171.43 | 87.43 | 93.15 |
| 2,048 | 55.41 | 68.58 | 135.45 | 181.12 | 200.62 | 124.15 | 115.44 |

**Table 13:** AtlasFold five-sample batching throughput at different maximum numbers of residues processed together.

| $L$ | Maximum residues processed together | | | | | | |
| --- | --- | --- | --- | --- | --- | --- | --- |
|  | 64 | 128 | 256 | 512 | 1,024 | 2,048 | 4,096 |
| 64 | 0.775 | 1.724 | 3.177 | 5.964 | 8.232 | 9.037 | 9.663 |
| 128 | – | 0.692 | 1.443 | 2.587 | 4.004 | 4.515 | 4.795 |
| 256 | – | – | 0.671 | 1.182 | 1.391 | 1.479 | 1.533 |
| 512 | – | – | – | 0.366 | 0.392 | 0.403 | 0.413 |

#### D.3 Complex inference profiling

The co-folding baselines are run one target at a time in this comparison. Baseline implementations and inference settings are specified in Section C.3.1. The comparison covers complexes with 256–2,048 total residues. Table 15 reports peak GPU memory during unbatched complex inference. Table 14 reports unbatched complex inference runtime. AtlasFold-M supports batched inference. The batching sweep uses five diffusion samples, 10 recycles, and 200 diffusion steps. Table 16 reports AtlasFold-M batching throughput.

**Table 14:** Unbatched inference runtime in seconds per complex.

| Length | AtlasFold-M | ESMFold2 | Boltz-1 | Protenix-v1 | AF-Multimer v2.3 | AlphaFold3 |
| --- | --- | --- | --- | --- | --- | --- |
| 256 | 6.52 | 3.76 | 14.64 | 12.94 | 15.09 | 3.00 |
| 384 | 7.77 | 6.24 | 22.56 | 12.44 | 25.39 | 6.43 |
| 512 | 11.04 | 9.89 | 35.85 | 13.40 | 33.84 | 6.43 |
| 640 | 15.19 | 14.78 | 61.19 | 18.15 | 46.17 | 12.00 |
| 768 | 20.08 | 19.94 | 85.39 | 21.48 | 62.91 | 12.00 |
| 896 | 26.24 | 27.90 | 100.76 | 28.04 | 75.54 | 20.45 |
| 1,024 | 33.24 | 39.17 | 158.44 | 34.96 | 90.59 | 20.45 |
| 1,280 | 52.46 | 61.93 | 219.46 | 55.62 | 135.54 | 32.95 |
| 1,536 | 76.64 | 92.67 | 381.11 | 82.43 | 192.95 | 47.91 |
| 1,792 | 107.32 | 132.85 | 473.02 | 117.74 | 254.86 | 91.50 |
| 2,048 | 145.90 | 181.12 | 719.34 | 171.73 | 346.78 | 91.50 |

**Table 15:**
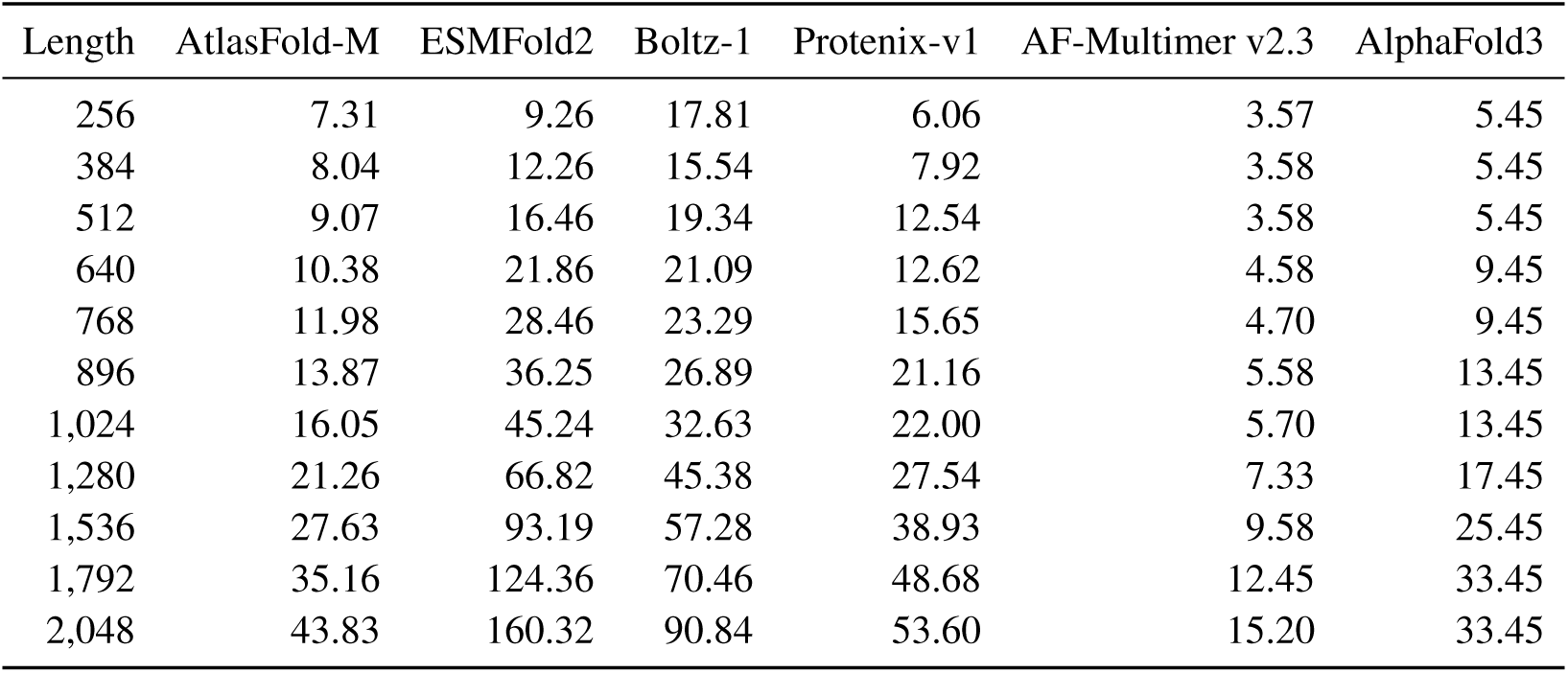
Peak GPU memory in GiB during unbatched inference.

**Table 16:** AtlasFold-M five-sample batching throughput at different maximum numbers of residues processed together.

| Length | 256 | 512 | 1,024 | 2,048 | 4,096 |
| --- | --- | --- | --- | --- | --- |
| 256 | 0.149 | 0.240 | 0.275 | 0.293 | 0.314 |
| 512 | – | 0.090 | 0.100 | 0.106 | 0.106 |

## Footnotes

1 https://github.com/SeonghwanSeo/atlasfold

2 20 steps for *L ≤* 512, 30 for 512 *< L ≤* 1,024, and 100 for *L >* 1,024, where *L* is sequence length.

3 https://portal.openfold.omsf.io/

4 https://github.com/SeonghwanSeo/esmfold-minimal

5 https://portal.openfold.omsf.io/

